# Structural mechanism governing radiationless energy transfer in *Renilla* bioluminescence

**DOI:** 10.64898/2026.09.01.748624

**Authors:** Daniel Pluskal, Martin Toul, Naina Verma, Anna Smrckova, Tadeja Gao, Tom Buryska, Tim de Martines, Jana Horackova, Marketa Stuchla, Veronika Novakova, Michal Nemergut, Yves L. Janin, Andrew DeMello, Stavros Stavrakis, Martin Vacha, Jiri Damborsky, David Bednar, Lenka Hernychova, Zbynek Prokop, Martin Marek

**Affiliations:** Loschmidt Laboratories, Department of Experimental Biology and RECETOX, Faculty of Science, Masaryk University, Kamenice 5, Bld. C13, 625 00 Brno, Czech Republic; International Clinical Research Center, St. Anne’s University Hospital Brno, Pekarska 53, 656 91 Brno, Czech Republic; Research Center for Applied Molecular Oncology, Masaryk Memorial Cancer Institute, Zluty kopec 7, 656 53 Brno, Czech Republic; Institute for Chemical and Bioengineering, ETH Zürich, 8093 Zürich, Switzerland; Structure et Instabilité des Génomes (StrInG), Muséum National d’Histoire Naturelle, INSERM, CNRS, Alliance Sorbonne Université, 75005 Paris, France; School of Materials and Chemical Technology, Institute of Science Tokyo, Ookayama 2-12-1, Meguro-ku, Tokyo 152-8552, Japan

**Keywords:** Bioluminescence, BRET, dipole-dipole coupling, *Renilla reniformis*, luciferase, resonance energy transfer

## Abstract

The nonradiative transport of electronic excitation from one chromophore to another, known as resonance energy transfer, lies at the root of photochemical processes in biology.^1,2^ Unlike photosynthesis, bioluminescence converts chemical energy into light through an enzymatic oxygenation of an energy-rich luciferin.^3–5^ In glowing cnidarians, the energy is relocated from an excited oxyluciferin to a fluorescent protein, shifting the colour and enhancing the quantum yield of a photogenic reaction.^6,7^ How protein-chromophore complexes assemble during this interplay in real space, and what this association entails for function, are unknown. Here, we report co-crystal structures of a 120-kilodalton energy-transfer complex from the luminescent soft coral *Renilla reniformis*. We find a heterotetrameric 2:2 assembly composed of two coelenteramide-loaded luciferases (RrLuc) docked at opposite sides of a head-to-tail dimer of green fluorescent protein (RrGFP). The edge-to-edge distance between donor and acceptor chromophores is below 3 nm, favouring the Förster-type radiationless energy transfer. Furthermore, RrGFP serves not only as a colour-switchable antenna and luminescence amplifier but also tunes the efficiency of luciferase catalysis by controlling its inherent dynamics. Our results provide detailed spatial information about intermolecular dipole-dipole coupling in *Renilla* bioluminescence, including the arrangement of donor-acceptor pairs that secure excited-state energy transfer with exquisite precision.

## Introduction

Radiationless transfer of electronic excitation between two molecules coupled by a dipole-dipole interaction lies at the root of photochemical processes in biology, notably photosynthesis^2,8^ and bioluminescence^6,7^. Central to oxygenic photosynthesis are the protein-pigment complexes, photosystems I and II, that convert the energy of sunlight into chemical potential energy^1,2^. In contrast to photosynthesis, bioluminescence converts chemical energy to visible light through a luciferase-catalysed or photoprotein-based oxygenation of a luciferin. Some marine animals, notably cnidarians, evolved an ability to relocate energy from an excited oxyluciferin to a partnering fluorescent protein via a dipole-dipole coupling mechanism, shifting the colour and enhancing the quantum yield of a photogenic reaction^7^. This phenomenon, known as bioluminescence resonance energy transfer (BRET), has significant importance to numerous areas of science and technology^9–12^. Yet the macromolecular complexes, securing the interplay between the electric dipoles of chromophore molecules in bioluminescence, have not been visualized at the atomic level.

Many glowing cnidarians emit green light, often with narrow emission spectra shaped by green fluorescent proteins (GFPs) that modify the primary bioluminescent emission. One of the most studied cnidarians is the sea pansy *Renilla reniformis*, which emits green bioluminescence (509 nm) as a defensive reaction to mechanical disturbance. *R*. *reniformis* produces green light through a complex process relying on a BRET process, which is the result of a coevolution of a cofactor-independent luciferase (RrLuc or RLuc) and a green fluorescent protein (RrGFP) in this organism^6,7^. When isolated, RrLuc luciferase catalyses the conversion of marine luciferin coelenterazine (CTZ) to coelenteramide (CEI) monoanion emitter, giving a flash of blue light (∼470–480 nm)^3^. However, in *R*. *reniformis* polyps, the bioluminescent system is localized to specialized light-emitting cells (photocytes), where RrLuc and RrGFP cooperate. Following dioxygen addition to the CTZ luciferin in the active site of RrLuc luciferase, the monooxygenation reaction converts CTZ luciferin to an excited CEI anion that donates energy to a *p*-hydroxybenzylidene-imidazolidone fluorophore (CRO) of antenna RrGFP protein^6,7^. The radiative transitions of the RrGFP fluorophore and the excited CEI anion undergo a dipole-dipole coupling of the Förster type, and the RrGFP protein consequently radiates the energy as its own fluorescence (509 nm)^7^. Several lines of evidence indicated that the shift of bioluminescence spectra via radiationless energy transfer to the antenna RrGFP protein involves assembly of a macromolecular complex through highly specific protein-protein interactions^13^. However, key questions remain unanswered: (*i*) how do individual protein-chromophore subunits physically interact with each other, and (*ii*) what is the mutual distance and orientation between the luminescent donor and the fluorescent acceptor that allows them to transfer energy via spectacularly efficient dipole-dipole coupling?

Here, we report atomistic structures of a 120-kilodalton BRET complex from *R*. *reniformis*, revealing a heterotetrameric architecture composed of two CEI oxyluciferin-loaded RrLuc luciferases that dock on opposite sides of the RrGFP dimer. The distance between the CEI donor and the CRO acceptor of RrGFP protein within the complex is below 3 nm, favouring the nonradiative Förster-type resonance energy transfer. Biochemistry experiments demonstrate that RrGFP not only acts as a colour-switchable transmitter, but also substantially fine-tunes the catalytic efficiency of RrLuc luciferase. Our results provide detailed spatial information about intermolecular dipole-dipole coupling in *Renilla* bioluminescence, and open exciting prospects for nature-inspired, artificial energy-transfer and optical processes.

## Results

### Reconstitution of the excited-state energy-transfer complex

To reconstitute the *Renilla* BRET complex, we first characterized full-length RrLuc and RrGFP proteins expressed in *Escherichia coli*. Luminescence of RrLuc and fluorescence of RrGFP show emission maxima at 470 nm and 509 nm, respectively (**Fig. 1a**). The maxima do not change when RrGFP is added to RrLuc at various concentrations. However, the intensity of RrLuc emission decreases with a concomitant increase in the total area under the RrGFP emission curve (**Fig. 1b**). The observed efficient transformation of the RrLuc emission spectrum into that of RrGFP confirms the occurrence of BRET between the two recombinant proteins, in agreement with experiments previously published by Ward and Cormier, performed on *Renilla* luciferase and fluorescent protein isolated directly from the source organism^13^.

**Fig. 1.**
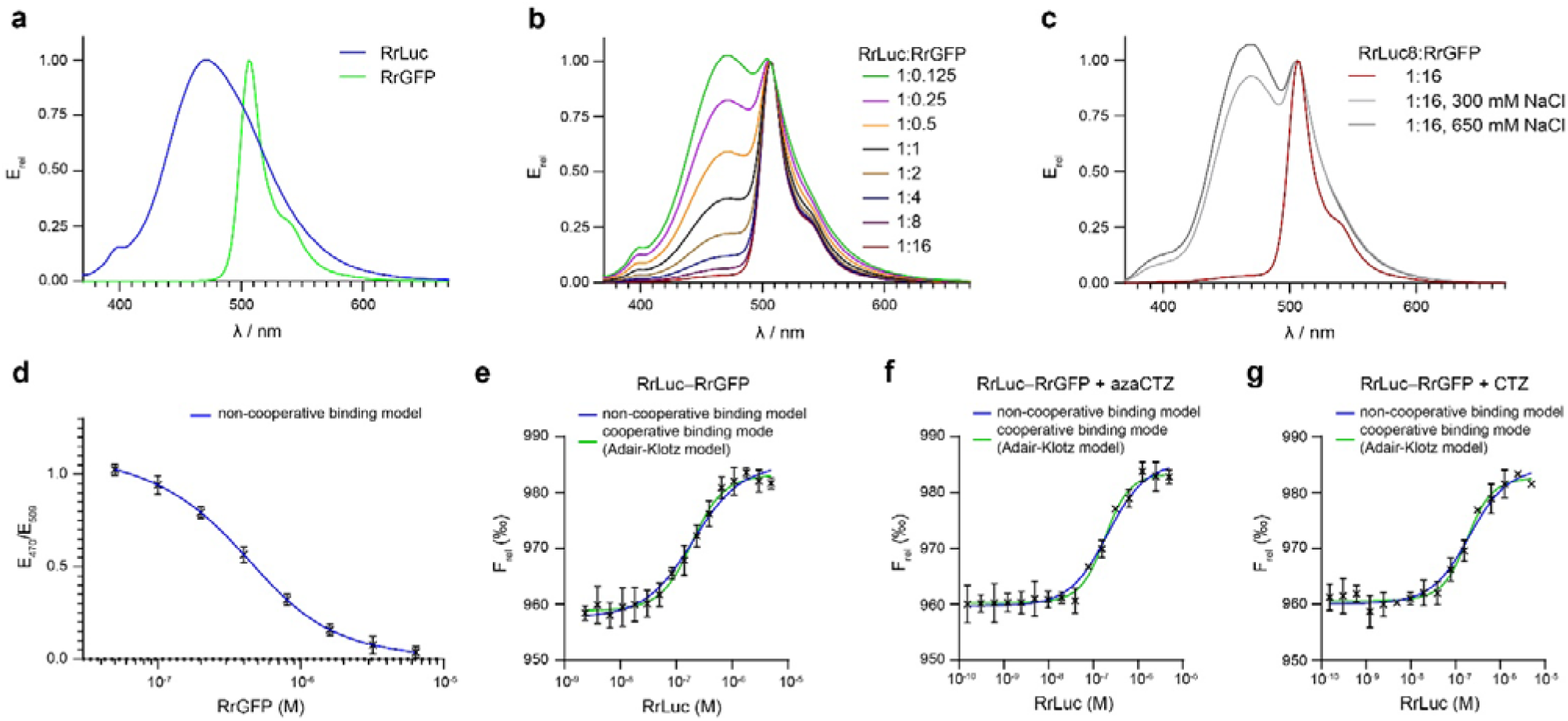
Characterisation of RrLuc□ RrGFP complexation. (**a**) Relative luminescence (CEI emitter) and fluorescence (CRO fluorophore) spectra of RrLuc and RrGFP, respectively. (**b**) Relative luminescence spectra of RrLuc in the presence of increased RrGFP concentrations, normalised based on emission at 509 nm. (**c**) Relative RrLuc luminescence in the presence of 16-fold molar excess of RrGFP at various ionic strengths, normalised based on emission at 509 nm. (**d**) Spectroluminometric determination of dissociation constant *K*_d_ of RrLuc□ RrGFP complexation, based on E_470_/E_509_ ratio at several RrGFP concentrations, fitted by noncooperative binding model. The solid blue line represents the best fit, and the error bars show the measurement S.D. (n = 3). (**e**□ **g**) Thermophoretic analysis of the interaction between RrLuc and RrGFP. The affinities were measured in the absence of CTZ substrate (e), in the presence of non-oxidizable analogue azaCTZ (f), or in the presence of CTZ substrate (g). Blue solid lines represent the best fits of the non-cooperative binding model, and the green solid lines represent the best fit of the cooperative Adair-Klotz model. The error bars show the measurement S.D. (n = 3). All measurements (**a-g**) were conducted in 5 mM KCl, 10 mM Tris buffer (pH = 7.5) at 25 °C.

To confirm that the observed BRET is a consequence of complexation between recombinant RrLuc and RrGFP rather than a result of random in-solution BRET, we performed analogous measurements at increased ionic strength, similar to those performed by Ward and Cormier on proteins isolated from *Renilla*^6^. Elevated salt concentrations substantially decrease the intensity of BRET (**Fig. 1c**). Although this result does not exclude the possibility of some random BRET and/or trivial energy-transfer events, it suggests that the BRET occurs via a physical contact between RrLuc and RrGFP mediated by an electrostatic interaction. The dissociation constant of RrLuc□RrGFP interaction (*K*_d_) was determined by the ratio of emission intensities (E_470_/E_509_) and provided the value 209 ± 23 nM (**Fig. 1d**).

### Positive cooperative binding

The effect of a substrate or product on binding affinity (*K*_d_) has been reported for some bacterial luciferases and *Clytia gregaria* photoprotein^14,15^. Therefore, this aspect was also addressed for RrLuc□ RrGFP complexation. Since the luminometry (**Fig. 1a-d**) requires CTZ-luciferin, we employed microscale thermophoresis and temperature-related intensity change (MST-TRIC) analysis instead, which allowed all measurements to be performed with or without CTZ/CEI molecules present (**Fig. 1e-g** and **Extended Data Table 1**). The RrLuc catalysis proceeds very rapidly (in seconds), making it challenging to capture the enzyme-substrate state. To overcome this issue, we used a non-oxidizable substrate analogue, azacoelenterazine (azaCTZ)^3^, which serves as a surrogate to probe the enzyme-substrate state.

Obtained values of apparent *K*_d_ (*K*_d,app_) agreed with the value retrieved from spectroluminometry (∼200 nM). The experiment revealed no significant effect of substrate or product presence on RrLuc−RrGFP interaction, yet, interestingly, showed a significant cooperativity in binding of RrLuc to RrGFP (Hill coefficient *n*_H_ ∼1.5). This contradicts the previously reported conclusion that only one RrLuc molecule binds per RrGFP dimer^6^. The overall RrLuc−RrGFP interaction appears to comprise two binding events, with the second binding event affinity being ∼10-fold stronger than the affinity of the first binding event (microscopic dissociation constants of first and second binding events were determined as 604 ± 54 nM and 62 ± 6 nM, respectively; **Extended Data Table 1**).

### Atomic resolution structures of the energy-transfer complex

To elucidate the structural organization of the *Renilla* BRET complex, full-length RrLuc and RrGFP proteins were co-expressed in *Escherichia coli* (**Fig. 2a**). The untagged RrGFP co-purified with C-terminally His-tagged RrLuc, further confirming the existence of a physical RrLuc−RrGFP complex. The purified complex showed vibrant green fluorescence and contained both RrLuc and RrGFP in near-stoichiometric amounts (**Fig. 2b**,**c**). Upon mixing with CTZ-luciferin, the complex yielded a flash of green light (509 nm), confirming that both catalytic and energy-transferring functions were retained. The CEI oxyluciferin-loaded RrLuc-RrGFP complex was then subjected to crystallization. We obtained greenish crystals under several conditions, from which some diffracted up to 2.3 Å resolution (**Fig. 2d**). The structures of the complex were solved by molecular replacement (**Extended Data Table 2**). The final electron density maps were sufficient to build all-atom protein models, including solvent-exposed and terminal regions that had previously been unresolved for free RrLuc or RrGFP^16^.

**Fig. 2.**
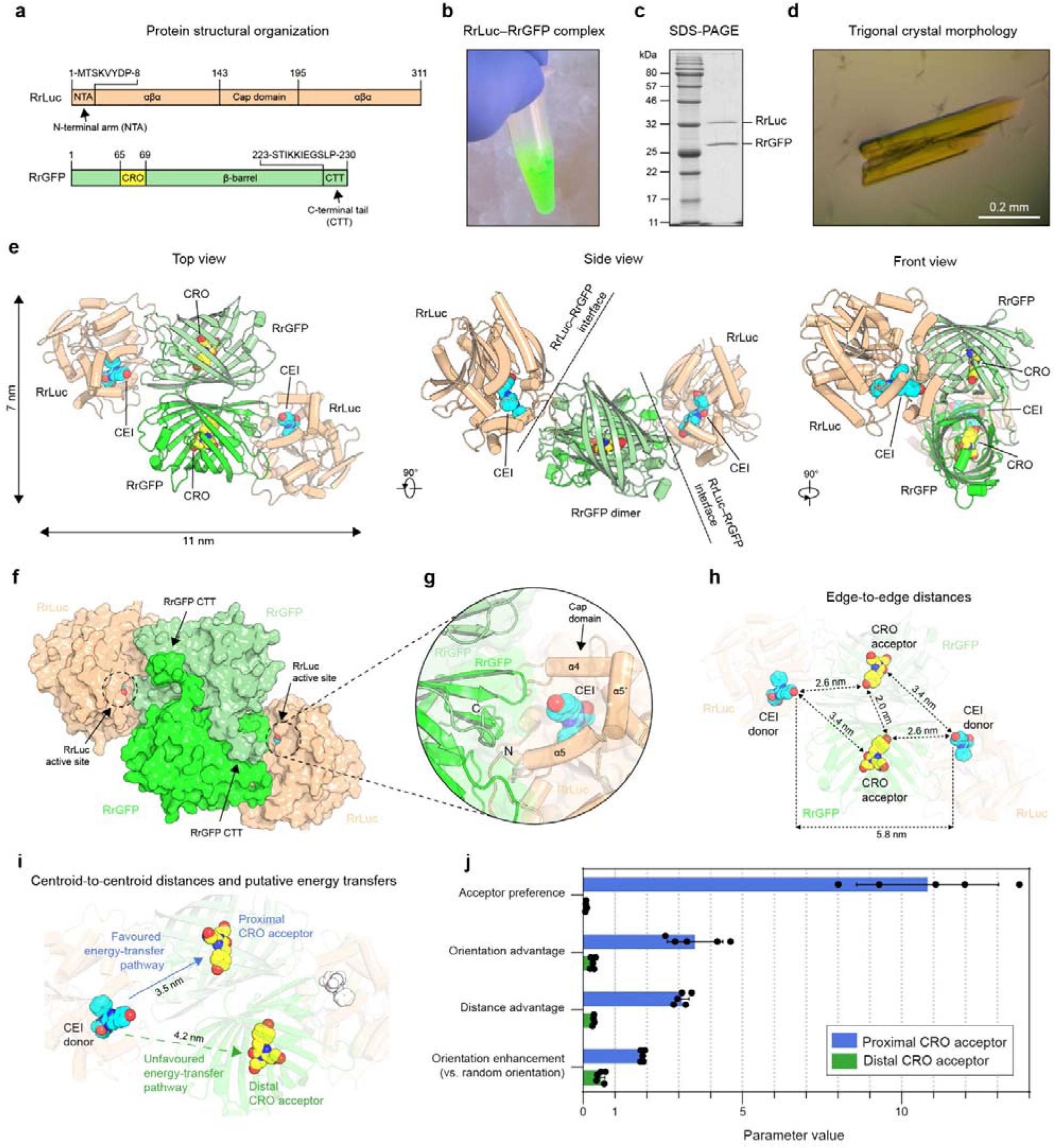
Structure of the *Renilla* BRET complex. (**a**) Schematic of the domain organisation of RrLuc and RrGFP proteins. (**b**) Green colour of the reconstituted RrLuc□ RrGFP complex. (**c**) SDS-PAGE analysis of RrLuc□ RrGFP complex. (**d**) Greenish trigonal crystal of CEI-bound RrLuc□RrGFP complex. (**e**) Overview of the CEI-bound RrLuc□ RrGFP complex shown in three orientations. RrGFP protomers are depicted in two shades of green, and RrLuc protomers are in beige. CRO fluorophores of RrGFP are depicted in yellow and CEI oxyluciferins in cyan. (**f**) Surface representation of the CEI-bound RrLuc□ RrGFP complex. The location of the RrLuc active site cavity with the bound CEI oxyluciferin is indicated with a dashed circle. The C-terminal tail (CTT) in both RrGFP protomers is indicated with a black arrow. (**g**) Magnified view of the RrLuc□RrGFP complexation with the depicted cap domain of RrLuc. CEI is shown in cyan. (**h**) Arrangement, orientation, and edge-to-edge distances between chromophores. The CRO of RrGFP is depicted in yellow and CEI oxyluciferins in cyan. (**i**) Centroid-to-centroid distances between CEI donor and CRO acceptors, with depicted two possible energy-transfer pathways. All structural figures (**e-i**) were generated from PDB entry: 8RZZ. (**j**) The proximal CRO is the preferred energy acceptor. The acceptor preference is calculated as a ratio of FRET efficiency factor *f*_DA_ for the particular CRO acceptor and the other CRO acceptor in the same RrGFP dimer. The acceptor preference can be decomposed into the orientation advantage, equal to the ratio of the FRET orientation factors κ^2^ of the proximal and distal acceptors (κ^2^_proximal_/κ^2^_distal_), and the distance advantage, calculated as the ratio of the sixth powers of the distal and proximal acceptor distances *r* (*r*^6^_distal_/r^6^_proximal_); the acceptor preference is the product of the orientation and distance advantages. The acceptor preference, orientation advantage, and distance advantage are defined relative to the proximal acceptor; for the distal acceptor, the corresponding values are the reciprocals of the defined ratios. Orientation enhancement is the fold-improvement in *f*_DA_ for a particular CRO acceptor over an equidistant acceptor with random orientation in space (κ^2^ = ⅔), highlighting the importance of the evolved CEI−CRO orientation to the FRET efficiency within the RrLuc−RrGFP complex.

The quaternary structure of *Renilla* BRET complex shows a heterotetrameric architecture (**Fig. 2e**,**f**). Two RrLuc luciferases dock at opposite sides of the RrGFP dimer, creating a 2:2 RrLuc□ RrGFP complex that is around 120 kDa in size. The structural observation concerning RrLuc−RrGFP complex stoichiometry thus agrees with our biophysical measurements (**Fig. 1**). The central part of the tetrameric complex is a head-to-tail homodimer with nearly perfect two-fold symmetry, formed by two RrGFP protomers that interact extensively with each other. Two single RrLuc modules are symmetrically attached on the facing sides of the RrGFP dimer, bringing the CEI oxyluciferin as close as possible to the CRO fluorophore of RrGFP, created by an autocatalytic cyclization of three amino acids (S66, Y67, and G68) inside the RrGFP protein. Due to the 2:2 stoichiometry, the heterotetrameric association therefore brings together two donor chromophores (CEI oxyluciferins) and two acceptor chromophores (CRO fluorophores).

For efficient resonance energy transfer to occur, the donor-acceptor separation must be below 10 nm (i.e., the acceptor and donor electric fields must meaningfully overlap)^17^. The CEI-loaded RrLuc−RrGFP co-complex fulfils this physical requirement: the edge-to-edge distance between the CEI oxyluciferin, bound in the catalytic pocket of RrLuc luciferase, and the CRO moiety found inside the RrGFP β-barrel, is ∼2.6 nm (**Fig. 2h**). This exceptionally short donor-acceptor separation falls well within the range required for near-quantitative Förster energy transfer. Such an arrangement suggests that the evolution of the BRET complex in *R. reniformis* optimized the donor-to-acceptor distance for highly efficient nonradiative Förster-type resonance energy transfer.

### Favoured energy transfer pathway

The structural analysis shows that for each RrLuc-bound CEI donor, there are two possible pathways for the Förster-type energy transfer: either via the CRO fluorophore of the RrGFP protomer in the main-interface interaction with the RrLuc binding the CEI oxyluciferin (proximal acceptor), or the CRO fluorophore of the other RrGFP protomer present in the 2:2 complex (distal acceptor). To calculate the likelihood of the respective pathways, we used the Förster resonant energy transfer (FRET) model^18^ (**Extended Data Table 3**). The energy transfer from any particular CEI donor to its proximal CRO acceptor was revealed to be ∼10 times more likely than that to the distal acceptor, both because of the closer distance and more favourable orientation. Furthermore, whereas the orientation of the proximal acceptor promotes efficient FRET, the distal acceptor is oriented unfavourably for energy transfer. The distal CRO fluorophore is constrained such that FRET to this acceptor is only about half as efficient as would be expected under the assumption of random transition dipole moment orientations. Consequently, each CEI donor exhibits a pronounced preference for transferring energy to its paired proximal CRO acceptor (**Fig. 2i**,**j**).

### The rewiring of RrGFP β-barrels for complexation

RrGFP, a β-barrel fold protein, exists as a compact head-to-tail homodimer^16^. Notably, the RrGFP structure previously determined by Loening and co-workers lacks electron density maps for several solvent-exposed loops and the C-terminal sequence (K_227_IEGSLP)^16^. Here, we crystallized and determined structures of full-length RrGFP in both apo- and RrLuc-bound states to reveal molecular changes essential for protein-protein complexation. The RrGFP fold consists of an 12-stranded β-barrel with an intra-barrel coaxial helix loop carrying the CRO fluorophore. The co-crystal structures reveal the reorganization of the RrGFP dimer, involving the β12 strand, L15 loop, and C-terminal tail (CTT), essential for harbouring two RrLuc luciferases (**Fig. 3a-e**). Structurally, the two RrGFP β-barrels are oriented upside down with respect to each other and tilted by ∼80° on the vertical axis, showing an assembly typical for many fluorescent proteins (**Extended Data Fig. 1**). PISA calculations on the full-length RrGFP showed that the homodimer interface area is ∼2,010□ Å^2^, which represents ∼17% of the total solvent-accessible surface area of the monomer (∼11,900□ Å^2^). There are 51 (chain A) and 53 (chain B) interfacing residues involved in the dimer interface, which is a significant portion of the 233-residue protein (**Fig. 3f**). The homodimeric interface is formed by up to 13 hydrogen bonds and numerous polar and nonpolar contacts.

**Fig. 3.**
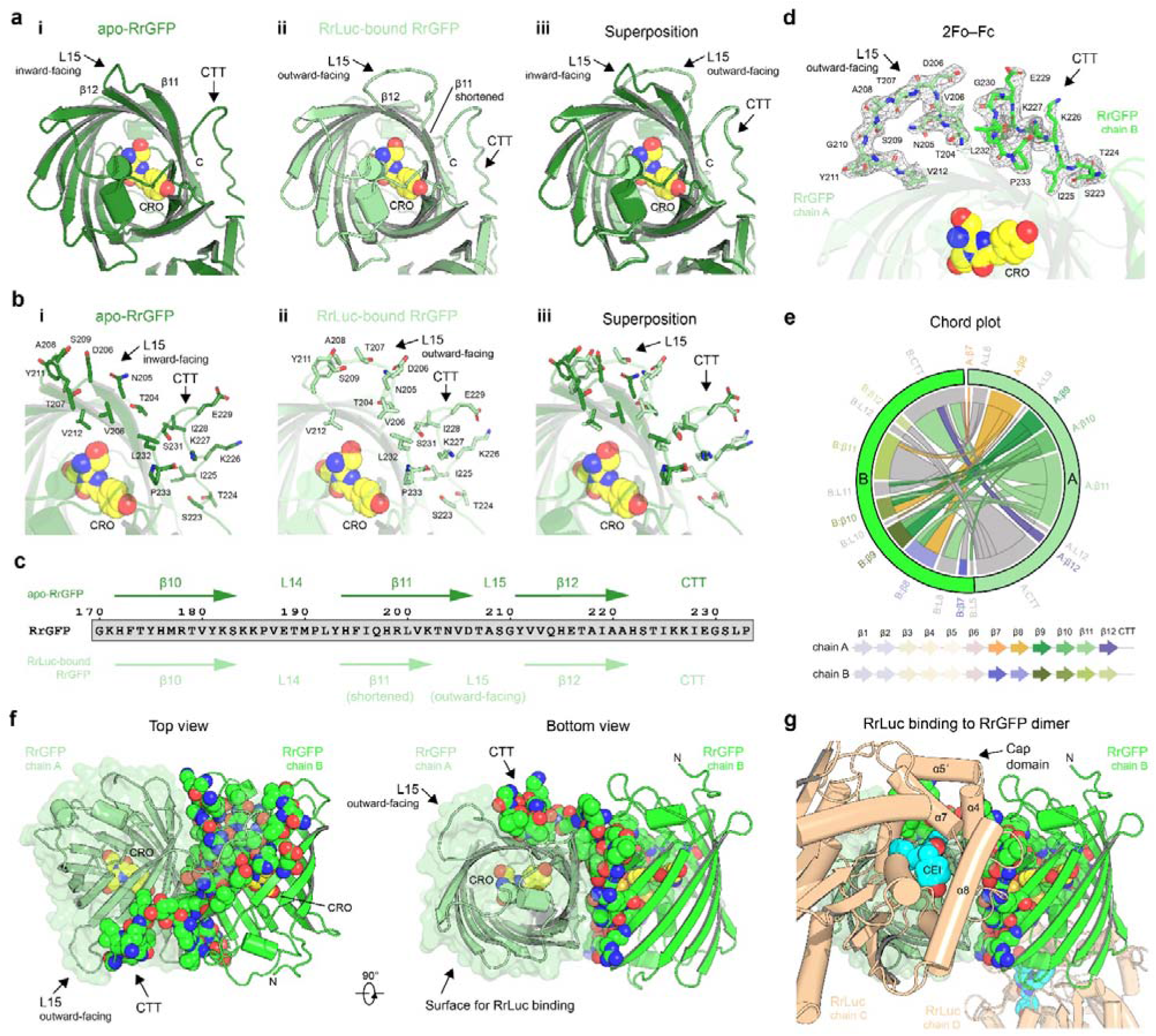
Structural rearrangements in RrGFP dimer required for complexation with RrLuc. (**a**,**b**) Comparison between apo RrGFP and RrLuc-bound RrGFP structures. Cartoon representations: (a) and close-up views (b) of apo RrGFP (i), RrLuc-bound RrGFP (ii), and superposed views (iii). Note structural rearrangements in the L15 loop and β11 strand. (**c**) Topology of secondary structure elements in apo RrGFP and RrLuc-bound RrGFP structures. (**d**) 2Fo-Fc electron density maps (contour level 1.2 σ) for the L15 loop (chain A) and CTT (chain B). The CRO fluorophore is shown as yellow spheres. (**e**) Chord plot showing the interactions between chains A and B in the RrGFP dimer at the secondary structure level, calculated and visualised by Protein Contact Atlas^20^. (**f**) Homodimer interface visualisation. Cartoon representation of chain A (pale green) and chain B (green) in the RrGFP homodimer. All residues in chain B involved in the dimer interface are shown as space-filling spheres. The CRO is shown as yellow spheres. (**g**) Close-up view of the RrLuc (beige) binding to the RrGFP homodimer. The CEI is shown as cyan spheres. All structural figures were generated from PDB entries: 8RZZ and 8S1V.

Despite the characteristic β-barrel fold, there are features distinguishing the RrGFP from its cnidarian counterparts (**Extended Data Fig. 1e**). Notably, the RrGFP CTT shows a unique sequence composition. In the homodimeric assembly, the CTT of one chain wraps around the surface of the twin β-barrel and *vice versa*, creating a nearly symmetrical dimerization interface. Notably, the mutual interconnection through the CTTs reinforces the stability of the RrGFP dimer and creates an interaction harbour for the docking of RrLuc modules.

The bottom part of RrGFP β-barrel serves as the docking platform for the RrLuc luciferase. Upon the RrLuc binding, the RrGFP β11 and β12 strands, together with the connecting L15 loop, undergo structural rearrangements to adopt better complementarity between the interacting protein surfaces (**Fig. 3f**,**g**). From an overall perspective, the analogous interaction harbour is created at the opposite side of the RrGFP dimer, ensuring the 2:2 RrLuc□ RrGFP complex stoichiometry.

### Noncovalent connection of submodules

The key to a radiationless BRET process is the connection between the energy-donating submodule, which is the CEI-bound RrLuc, and the energy-accepting RrGFP antenna. The orientation of β-barrels in the RrGFP submodule ensures no steric restrictions for the simultaneous binding of two RrLuc modules, each from one side of the RrGFP dimer. Thus, there are two identical RrGFP□ RrLuc interfaces in the resulting BRET complex, providing numerous polar and nonpolar contacts (**Fig. 3g**; **Extended Data Fig. 2**; **Extended Data Table 4**).

Structurally, the RrLuc exhibits a characteristic α/β-hydrolase fold. The enzyme core is composed of a central 8-stranded β-sheet surrounded by several α-helices. In the BRET complex, the RrLuc docks at the bottom part of one RrGFP chain. While all crystal structures of free RrLuc determined so far showed substantial conformational freedom of its N-terminal arm (NTA)^3,16,19^, this is not the case when it interacts with RrGFP. The structures of RrLuc□ RrGFP complex reveal that the RrLuc NTA (M_1_TSKVYDP) folds back towards the cap domain, and inserts into a groove shaped at the bottom of RrGFP (**Extended Data Fig. 2a**). Another striking feature seen in the BRET complex is the conformationally restricted cap domain of RrLuc. Our previous work highlighted that the cap domain of RrLuc explores a large conformational space, enabling multiple active-site cavity opening states^3^. The structural elements responsible for this cap domain malleability are the α4 helix, the L9 loop, and the L16 loop. Here, we show that the CEI-bound cap domain of RrLuc, when complexed with RrGFP, adopts a minimally open state. The RrLuc□RrGFP interface is formed by numerous hydrogen bonds and salt bridges (**Extended Data Fig. 2c-d**). Mutagenesis experiments of residues involved in the protein-protein interface confirmed the relevance of co-crystal structures (**Extended Data Fig. 2e,f**), as substituting these residues yielded no RrLuc–RrGFP complex formation in nearly all mutants.

### Allosteric signal explains cooperativity

Our biochemistry experiments revealed that the binding affinity of the second RrLuc is ∼10-fold stronger than the affinity of the first one, suggesting cooperative behaviour. The structures of *Renilla* BRET complex show 2:2 stoichiometry, with two identical RrLuc-harbouring sites symmetrically located on opposite sides of the RrGFP dimer. Careful inspection revealed that while one RrLuc molecule predominantly interacts with its proximal chain of RrGFP with a buried interface area of ∼1,041□ Å^2^, the CTT of the distal RrGFP molecule protrudes into its proximity and interacts with the same RrLuc molecule too, with a marginal contribution of buried area (∼37□ Å^2^). Although this contribution to the interfacing area is low, it helps the other GFP chain to restructure for the RrLuc binding (**Fig. 3f,g**).

Specifically, structures show that the binding of RrLuc triggers structural rearrangements in the RrGFP dimer, suggesting that the emergence of the first RrLuc binding site on one side of the RrGFP dimer can be transmitted via an allosteric signalling to the second binding site. We infer that the allosteric signal is transferred over a long distance (∼6.5 nm) through the C-terminal part of RrGFP (aa 203-233), encompassing the L15 loop, long β12 strand with adjacent CTT, which connect the two RrLuc binding sites in the homodimeric assembly (**Extended Data Fig. 1d,e**). Consequently, the pre-shaped second binding interface can harbour the second RrLuc enzyme with higher binding affinity. The structural transitions found here can explain the observed positive cooperative behaviour during the heterotetrameric BRET complex assembly (**Fig. 1**, **Extended Data Table 1**).

### Zooming in on the donor-acceptor pair

The mutual orientation between the CEI donor and CRO acceptor is shown in **Fig. 4a**. The 4-(p-hydroxybenzylidene)imidazolidin-5-one CRO fluorophore in RrGFP is generated from the autocatalytic cyclization and oxidation of three consecutive amino acids (S_66_YG). The CRO fluorophore is covalently linked and rigidly encapsulated inside the β-barrel, making numerous contacts and hydrogen bonds with adjacent residues and water molecules (**Fig. 4b-e**).

**Fig. 4.**
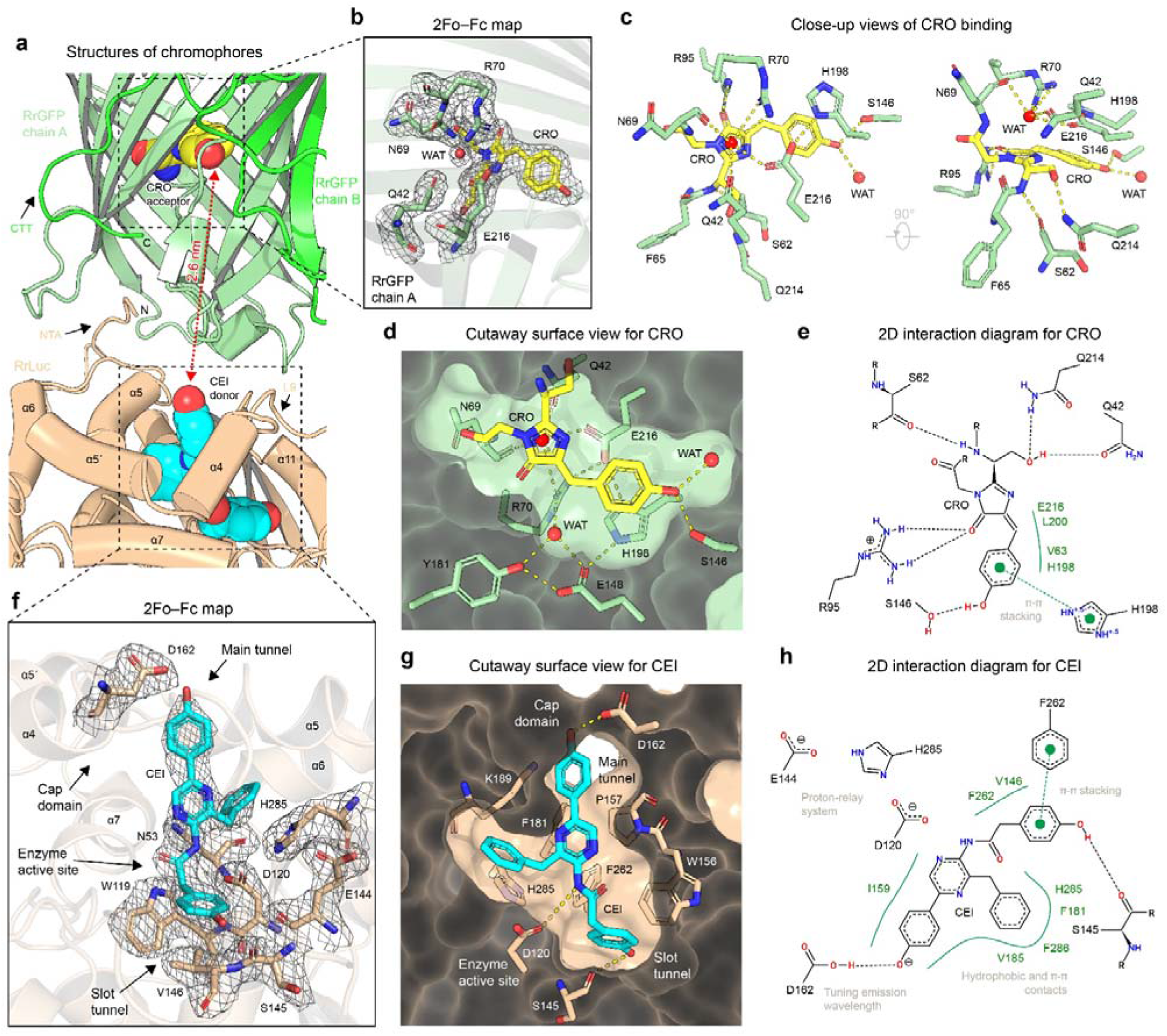
Molecular structures and orientations of energy donor and acceptor chromophores. (**a**) Cartoon representation of RrLuc□RrGFP complexation. The CEI (energy donor) is shown as cyan spheres, and the CRO (energy acceptor) is shown as yellow spheres. (**b**) 2Fo-Fc electron density maps (contour level 1.2 σ) for the CRO (RrGFP). (**c**) Close-up views of CRO within the RrGFP. Water molecules are shown as red spheres. Key molecular contacts are shown as dashed yellow lines. (**d**) Cut-away surface view of CRO binding pocket. (**e**) Two-dimensional diagram depicting the CRO interactions, generated by PoseView^21^. (**f**) 2Fo-Fc electron density maps (contour level 1.2 σ) for the CEI (cyan) bound in RrLuc (beige). (**g**) Cut-away surface view of CEI-binding pocket in RrLuc. Key molecular contacts are shown as dashed yellow lines. (**h**) Two-dimensional diagram depicting the CEI□ RrLuc interactions, generated by PoseView.^2^ All structural figures were generated from PDB entry: 8RZZ.

Inspections of the electron density maps unambiguously revealed one CEI molecule bound in the catalytic pocket of both RrLuc luciferases (**Fig. 4f**). Due to the 2:2 RrLuc□ RrGFP stoichiometry, there are in total two CEI oxyluciferins in the BRET complex, separated by an edge-to-edge distance of ∼5.8 nm (**Fig. 2h**). The CEI, or N-[3-benzyl-5-(4-hydroxyphenyl)pyrazin-2-yl]-2-(4-hydroxyphenyl)acetamide, adopts a characteristic Y-shaped conformation when bound in the luciferase pocket (**Fig. 4f-h**). The amide nitrogen is in the vicinity of the carboxyl side chain of D120 (∼3.8 Å), favouring the protonation of CEI through a conserved proton-relay system^3^. The R^1^ 2-(4-hydroxyphenyl) substituent is deeply inserted in the slot tunnel, engaging in π-π stacking with F262 (∼3.7 Å). Moreover, its terminal hydroxyl moiety is hydrogen-bonded with the backbone carbonyl of S145 (∼ 2.6 Å). The other two substituents, the R^2^ 5-(4-hydroxyphenyl) and R^3^ 3-benzyl, together with the central pyrazin-2-yl moiety, reside in the main tunnel and form multiple contacts, most of which are hydrophobic or aromatic. The R^3^ 3-benzyl group makes hydrophobic contacts with F180, F181, V185, H285, F261, and F286. The capping hydroxyl group of the R^2^ 5-(4-hydroxyphenyl) substituent makes a hydrogen bond with the carboxyl side chain of D162 (∼2.6 Å), an aspartate residue located at the rim of the catalytic cavity (helix α4).

The binding mode of CEI oxyluciferin in the RrLuc catalytic pocket ensures that its two hydrophilic hydroxyl groups are exposed to bulk solvent. The R^1^ 2-(4-hydroxyphenyl) substituent through the slot tunnel, while the R^2^ 5-(4-hydroxyphenyl) via the main tunnel entrance (**Fig. 4g**). The latter moiety is oriented towards the RrLuc□ RrGFP interaction interface, shielded by the β8-β9 loop of RrGFP. Although there are no visible direct contacts between the CEI R^2^ 5-(4-hydroxyphenyl) substituent and the RrGFP β8-β9 loop residues (Q168 and T169), we hypothesize that some interactions can occur, for example, via water-mediated hydrogen bonds.

### Control of the luciferase dynamics

Previous crystallographic studies of RrLuc luciferase revealed its high structural flexibility, predominantly spanning an N-terminal arm (NTA) and a helical cap domain (**Fig. 5a**)^3,16,19^. The uncontrolled opening of the RrLuc catalytic pocket, which can accommodate two or more luciferin molecules at the same time, has been linked with enzyme inhibition^3^. The structure of the CEI-loaded RrLuc□ RrGFP complex revealed that the RrGFP binding restricts the conformational freedom of the RrLuc catalytic pocket (**Fig. 5b**). The complexation with RrGFP shields the most flexible cap domain, as well as the NTA that inserts into a groove shaped at the RrLuc□ RrGFP interface (**Fig. 5c-e**).

**Fig. 5.**
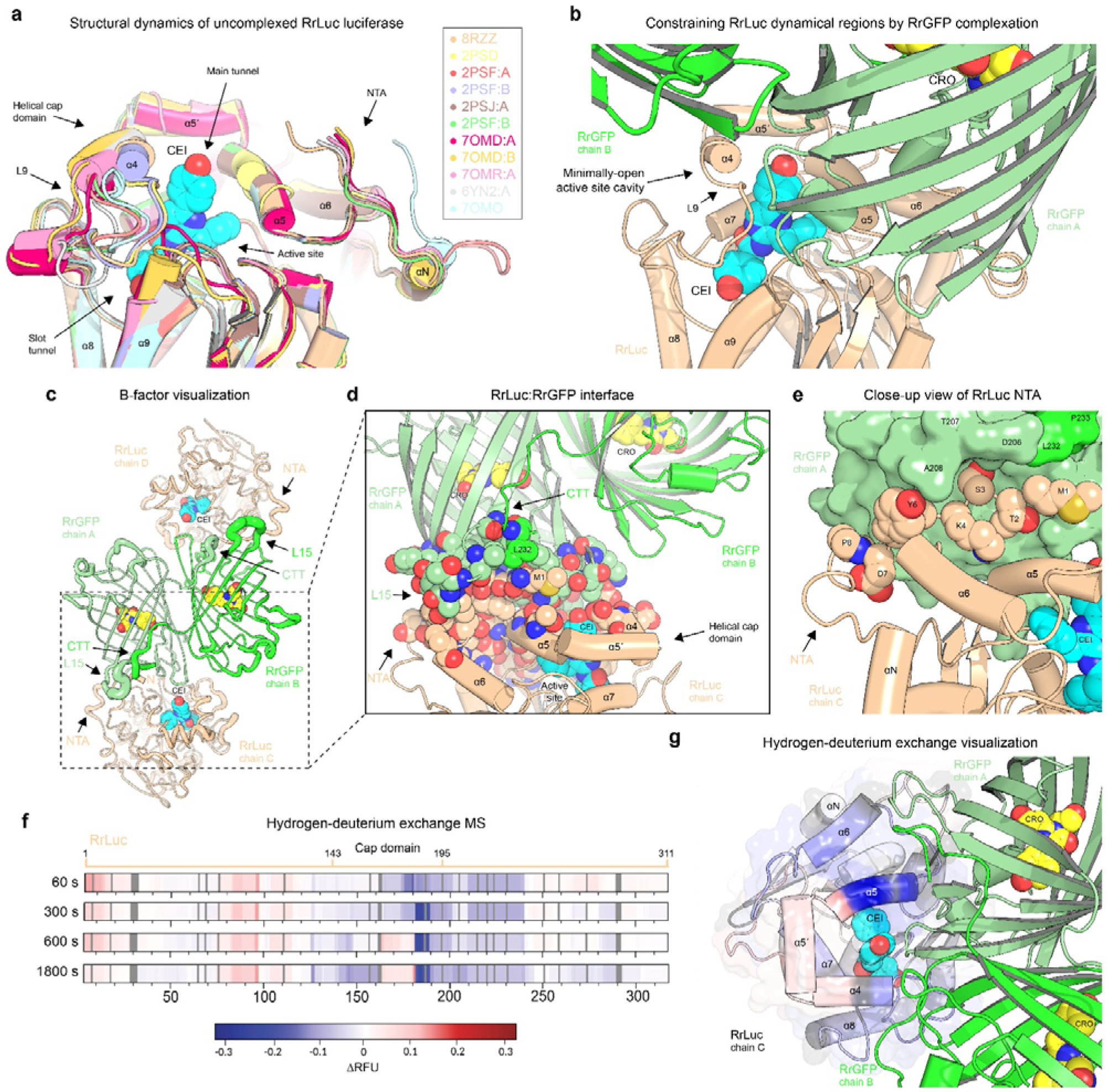
Controlling the RrLuc dynamics through a protein-protein complexation. (**a**) Superposed crystal structures of RrLuc luciferase, highlighting its structural flexibility. Major structural motions are seen in the cap domain and N-terminal arm (NTA). All PDB ID codes are listed. (**b**) Snapshot showing how the complexation with the RrGFP (green) restricts conformational malleability of RrLuc luciferase (beige). (**c**) B-factor putty representation of RrGFP□ RrLuc heterotetramer. (**d**) Visualisation of the RrGFP□RrLuc interface. All residues involved in the interface are shown as space-filling spheres. The CRO is shown as yellow spheres, and the CEI is shown as cyan spheres. (**e**) Zoomed view of the RrLuc NTA inserted in a groove shaped on the RrGFP surface. (**f**) HDX-MS analysis of unbound and RrGFP-bound RrLuc. Differences in relative fractional uptake were calculated as ΔRFU = RFU_RrGFP−bound_ – RFU_unbound_ and plotted along the RrLuc sequence after 60, 300, 600, and 1,800 s of deuterium exposure. Blue regions indicate protection upon complex formation (decreased amide hydrogen (H) to deuterium (D) exchange relative to the unbound RrLuc protein), while red regions indicate increased exchange upon binding. Proline residues, non-exchangeable in HDX-MS, and unresolved residues are indicated by dark and light grey rectangles, respectively. (**g**). Mapping of ΔRFU values onto the RrLuc structure within the BRET complex after 1,800 s of D_2_O labelling. The RrGFP dimer is shown in green cartoon representation, while RrLuc is coloured according to the ΔRFU scale. The CEI oxyluciferin and CRO fluorophore (energy acceptor) are shown as cyan and yellow spheres, respectively. All structural figures (**b-g**) were generated from PDB entry: 8RZZ.

To validate the RrLuc□RrGFP complexation observed in a crystal lattice, we conducted hydrogen-deuterium exchange coupled with mass spectrometry (HDX-MS) experiments. Peptide identification yielded 359 peptides providing 99.6% sequence coverage of the RrLuc. After manual quality control, 317 peptides were retained for HDX analysis, corresponding to 96.5% sequence coverage. The difference results from the exclusion of peptides with insufficient signal quality or unreliable isotopic envelopes. The labelling time points of 60, 300, 600, and 1800 s revealed structural elements in RrLuc, whose dynamics are controlled by protein-protein complexation (**Fig. 5f**; **Extended Data Fig. 3**).

To visualize binding-induced differences in deuterium uptake, we calculated ΔRFU (RFU_RrGFP-bound_ − RFU_unbound_), plotted the values along the RrLuc sequence (**Fig. 5f**), and mapped them onto the RrLuc–RrGFP complex structure (**Fig. 5g**). Complexation predominantly decreased deuterium uptake in multiple segments spanning D120–K300, consistent with local protection and/or reduced backbone dynamics, whereas the A164–F181 region showed increased uptake. The protected V141–E161 and V182–K189 segments are located at or near the RrGFP-binding interface. In contrast, the increased uptake within A164–F181 may reflect a local conformational rearrangement associated with increased solvent accessibility and/or backbone dynamics adjacent to the interface. Protection of the V141–E161 segment is consistent with the loss of complex formation observed for the D158A/E160A/E161A mutant, although only D158 directly contacts RrGFP in the crystal structure (**Extended Data Fig. 2e**). K189 was also identified as important for complex formation, while protection of the V182–K189 segment further supports the involvement of this region in the interaction interface. Overall, the HDX-MS data support the RrLuc–RrGFP interface observed in crystal structures and indicate that the RrGFP modulates the local conformational dynamics of RrLuc.

### Molecular simulations confirm constrained dynamics

The dynamics of CEI unbinding were computationally probed by the generation of Markov state models (MSM) from a total of 20 µs of adaptive sampling molecular dynamics (MD). Two macrostates were identified for the RrLuc–RrGFP complex, corresponding to a “bound” and “unbound” state of CEI (**Extended Data Fig. 4a-c**). The “bound” state represented most snapshots for the RrLuc–RrGFP complex, with an equilibrium probability of over 99.99%, indicating negligible oxyluciferin product release during the simulated period (**Extended Data Fig. 4d**). This outcome is supported by unbinding kinetics analysis resulting in a negative ΔG of the binding (**Extended Data Fig. 4f**). Surprisingly, previous comparable simulations of uncomplexed RrLuc demonstrated strikingly different characteristics. Here, the equilibrium probability of the “bound” state, defined as the combination of “bound” and “transit” states, was 57.2%^22^ (**Extended Data Fig. 4d**). Moreover, product release was energetically favoured in these simulations. Thus, protein simulations indicate a substantial shift in structural dynamics leading to a reduction in CEI product release when RrLuc is complexed with RrGFP.

To shed light on these altered dynamics, we studied the flexibility of RrLuc between an uncomplexed pose in comparison to its RrGFP-complexed pose. Uncomplexed RrLuc presents with generally higher flexibility, although peak profiles are comparable between both systems (**Extended Data Fig. 4e**). However, a considerably lower flexibility can be observed for the RrLuc–GFP complex in the region between the residues 145 and 180, corresponding to the cap domain of RrLuc^16^. Previous studies have demonstrated a role of this domain in substrate binding and product release mechanisms, with its flexibility crucial for adequate ligand transport in enzymes sharing the same fold^23^. Thus, the noticeably lower flexibility in this region observed in the RrLuc–GFP complex plausibly alters the dynamics of the enzyme-product complex, explaining the greatly reduced product release observed. This region likely has reduced flexibility due to its interaction with RrGFP in the complex, which acts as a physical barrier for the product through obstruction of the entrance tunnel (**Fig. 2g**). In summary, the interaction between RrLuc and RrGFP in the complexed form revealed reduced flexibility in the cap domain, which presents a notable obstacle in substrate binding and product unbinding, but suppresses unproductive binding and favours chemical catalysis.

### Protein-protein complexation empowers luciferase catalysis

To assess how complexation with RrGFP modulates the catalytic function of RrLuc, we performed a comprehensive kinetic analysis integrating steady-state and pre-steady-state measurements (**Fig. 6**; **Extended Data Table 5** and **Extended Data Fig. 5**). The data were globally interpreted within an induced-fit reaction framework (**Fig. 6a**), enabling a unified mechanistic description of CTZ binding and conversion by the RrGFP-bound RrLuc and inhibition of the reaction by a non-oxidizable CTZ analogue (azaCTZ^3^).

**Fig. 6.**
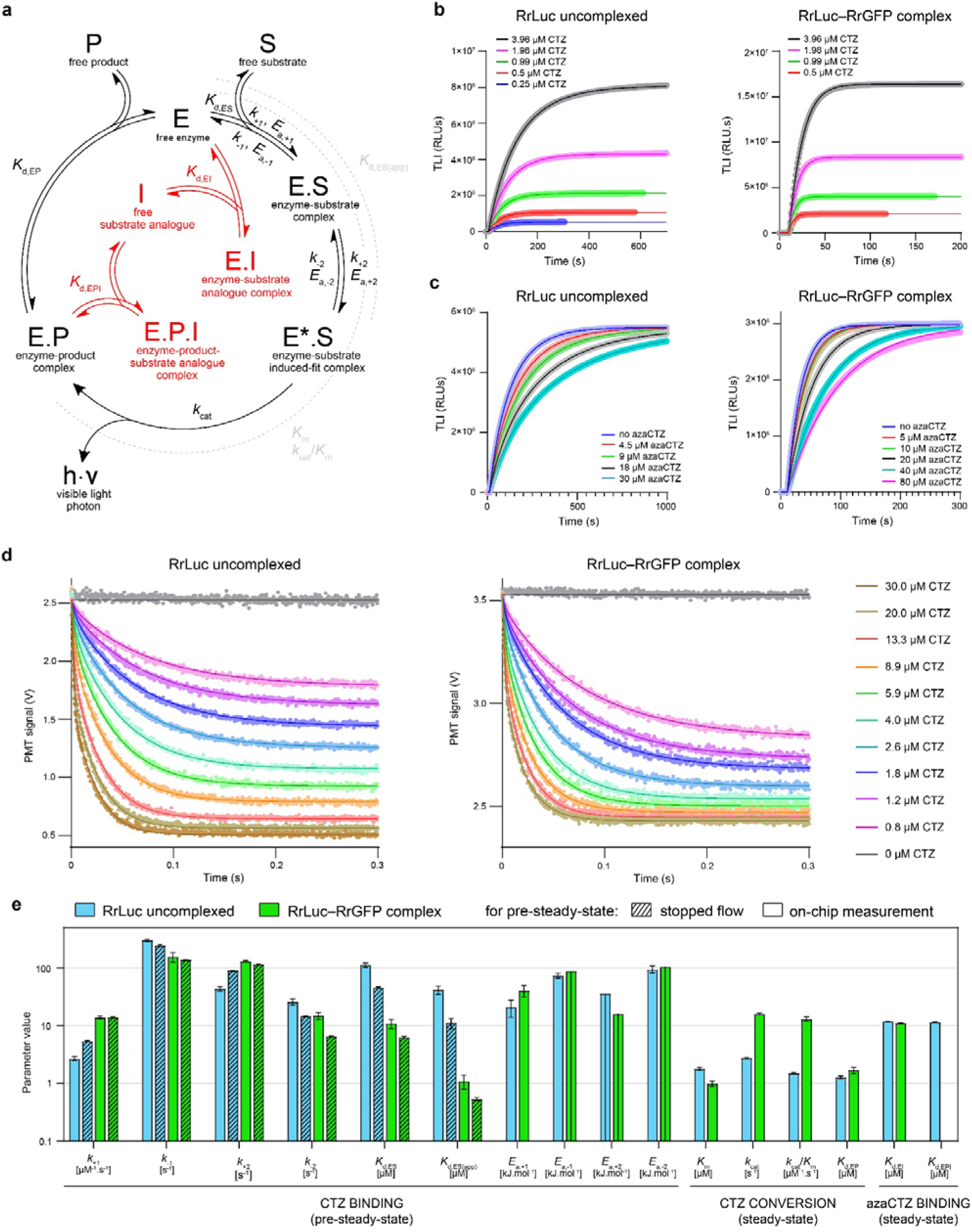
The complexation with RrGFP boosts the catalytic efficiency of RrLuc luciferase. **(a)** Overall model of CTZ conversion by RrLuc or RrLuc-RrGFP complex, indicating the assignment of all evaluated kinetic and thermodynamic parameters to individual reaction steps. Parameters describing grouped steps are shown in grey with dashed arcs. Black denotes the native catalytic cycle; red denotes the inhibition by azacoelenterazine (azaCTZ), a substrate analogue. *k*_+*n*_ = forward rate constant of reaction step *n*, *k*_–*n*_ = reverse rate constant of reaction step *n*, *K*_d,*X*_ = dissociation constant of species *X*, *K*_d,*X*(app)_ = apparent dissociation constant of species *X*, *E*_a,±*n*_ = activation energy of reaction step *n* (forward and reverse, respectively), *k*_cat_ = turnover number, *K*_m_ = Michaelis constant. **(b)** Total luminescence intensity (TLI) during coelenterazine conversion by RrLuc or RrLuc−RrGFP complex at varying CTZ concentrations. **(c)** TLI during conversion of 2 µM CTZ by RrLuc or RrLuc−RrGFP complex in the presence of increasing concentrations of azaCTZ (non-oxidisable substrate analogue). All CTZ conversion experiments were performed in a 5 mM KCl, 10 mM Tris buffer (pH = 7.5), in a triplicate. (**d**) Stopped-flow traces of protein tryptophan fluorescence during pre-steady-state CTZ binding to RrLuc or RrLuc−RrGFP complex collected at varying CTZ concentrations. The stopped-flow experiment was performed in 5 mM KCl and 10 mM Tris buffer (pH = 7.5), with each fitted trace being the average of 10-15 rapid-mixing experiments. For panels (**b**-**d**), symbols represent experimental data and solid lines indicate the best fits. (**e**) Summary of kinetic and thermodynamic parameters for RrLuc and the RrLuc−RrGFP complex derived from the reaction model depicted in panel (**a**). Parameters in the “CTZ binding” section were evaluated from stopped-flow data (panel **d**) and additional pre-steady-state on-chip measurements (**Extended Data** Fig. 5). Parameters in “CTZ conversion” and “azaCTZ binding” sections were derived from steady-state data shown in panels (**b**) and (**c**), respectively. Error bars represent the standard error (S.E.) of the fit. Confidence contour analysis revealed that the data did not define confident lower limits for three of the activation energies (*E*_a,-1_, *E*_a,+2_, *E*_a,-2_), so only upper limits could be determined.

Under steady-state conditions, complex formation with RrGFP substantially enhances the catalytic performance of RrLuc. The uncomplexed RrLuc exhibited catalytic parameters in agreement with previously reported values^3,19,22^, with minor deviations attributable to buffer composition optimized for RrLuc–RrGFP complexation. The RrLuc–RrGFP complex exhibits a ∼6-fold increase in turnover number (*k*_cat_) and a concomitant reduction in the Michaelis constant (*K*_m_). Together, these changes result in a ∼9-fold increase in catalytic efficiency (*k*_cat_*/K*_m_) (**Fig. 6e**; **Extended Data Table 5**). In addition, the RrLuc–RrGFP complex shows an increase in total luminescence intensity across all substrate concentrations (**Fig. 6b**), consistent with increased substrate utilization efficiency and apparent bioluminescence quantum yield^13^.

Pre-steady-state experiments (**Fig. 6d** and **Extended Data Fig. 5a,b**) confirm that CTZ binding to RrLuc–RrGFP complex proceeds via a two-step induced-fit mechanism we previously established for free RrLuc^3,19^, involving (i) rapid formation of an encounter complex E.S, followed by (ii) a conformational transition to the catalytically competent state E*.S. Here, we extended this analysis with temperature-dependent kinetics and thermodynamics measurements using an in-house rapid-mixing microfluidic chip^24^ and compared the results with a conventional stopped-flow experiment, yielding a good consistency across both methods. The quantitative analysis further shows that the improved catalytic efficiency of RrLuc–RrGFP is partly attributed also to enhanced substrate recognition and binding. The complexation of RrLuc and RrGFP accelerates both forward CTZ-binding steps: the bimolecular association rate constant (*k*_+1_) increases by ∼3–5-fold, while the conformational transition rate constant (*k*_+2_) is enhanced by ∼2–3-fold. In contrast, the reverse rates (*k*_-1_ and *k*_-2_) are reduced. Mechanistically, this indicates that RrGFP facilitates initial substrate capture by RrLuc and promotes the conformational locking of the enzyme-substrate complex, shifting the binding equilibrium toward catalytically productive states. These kinetic changes translate into 1-2 orders of magnitude increase in overall substrate affinity (reduced *K*_d,ES_ and *K*_d,ES(app)_ in **Fig. 6e** and **Extended Data Table 5**).

### Positive implications for function and mechanism

Additional insights into energetic barriers of the catalytic cycle and their differences for free and RrGFP-complexed RrLuc are provided by the activation energies derived from temperature-dependent pre-steady-state measurements (**Extended Data Table 5**). Notably, the activation energy of the initial association step (*E*_a,+1_) increases upon RrLuc complexation with RrGFP. In contrast, the activation barrier for the conformational transition (*E*_a,+2_) is reduced. Together with the kinetic observations, these changes indicate a redistribution of barriers, with substrate entry more tightly gated while progression towards the productive conformation is favoured, in agreement with the presented MD simulations. Consistent with these kinetic and energetic changes, the RrGFP-complexed luciferase shows reduced sensitivity to product inhibition. We previously described that unbound RrLuc is prone to forming an inhibitory enzyme–product (E.P) complex^3,19^. Although the equilibrium dissociation constant of this complex (*K*_d,EP_) remains in a similar micromolar range for both free and RrGFP-bound luciferase, the stabilization of substrate-bound enzyme states and their faster depletion through increased turnover effectively reduce the steady-state population of E.P complex and shift the reaction balance in favour of forward catalytic flux (**Extended Data Table 5**).

Inhibition experiments using azaCTZ further support this shift in pathway selectivity (**Fig. 6c**). AzaCTZ acts as a non-reactive substrate analogue that competes for the active site while also interacting with post-catalytic states^3^, consistent with a mixed-type inhibition model depicted in **Fig. 6a**. Both free and RrGFP-bound RrLuc exhibit similar affinity for E.I complex formation (*K*_d,EI_), indicating comparable competitive binding of azaCTZ to the active site. However, only free RrLuc forms an additional E.P.I complex (*K*_d,EPI_), whereas no such state is detected for the RrGFP-bound enzyme (**Fig. 6e** and **Extended Data Table 5**). This supports the notion that complexation with RrGFP restricts the luciferase conformational heterogeneity, suppressing the formation of off-pathway inhibitory states, effectively simplifying the reaction landscape, and improving the overall catalytic effectivity.

Altogether, the combined kinetic and thermodynamic analysis demonstrates that RrGFP acts as a functional modulator of RrLuc, redistributing flux within the kinetic framework. Acceleration of substrate binding and stabilisation of the induced-fit conformation, together with enhanced catalytic turnover, increased substrate utilization efficiency, and suppression of inhibitory states, collectively give rise to a more efficient, better-coordinated catalytic cycle and ultimately lead to much higher luminescence output.

## Discussion

Because of the extremely high efficiency of chemical energy conversion into light, there is a huge interest in harnessing bioluminescence for ultrasensitive optical bioassays, biosensor technologies, and even zero-electricity lighting sources. However, our contemporary knowledge of bioluminescence at a molecular level is still limited. This is particularly the case for interaction between electric dipoles of chromophore molecules in bioluminescence, the phenomenon known as BRET. In BRET, the energy is transferred from an excited donor molecule to a nearby unexcited acceptor molecule via intermolecular long-range, dipole-dipole coupling, without a need for physical contact between molecules as long as their distance does not exceed 10 nm^17^. However, in the homogeneous micromolar solutions of RrLuc and RrGFP, where BRET was observed, the distance exceeds the required length by an order of magnitude. Therefore, Ward and Cormier suggested that there should be physical contact between RrLuc and RrGFP^6^, but all attempts to capture the macromolecular BRET complex have failed so far. To address this challenge, here we biochemically and structurally characterized the BRET complex from the sea pansy *R*. *reniformis*, illuminating the molecular interplay between oxyluciferin-loaded RrLuc luciferase and RrGFP fluorescent protein that drives Förster-type radiationless energy transfer. A consequence of macromolecular complexation is that the RrLuc luciferase catalyses the photogenic reaction more efficiently, as evidenced by higher substrate affinity, reduced product inhibition, enhanced catalytic efficiency, and a higher total luminescence output.

Concerning the macromolecular complex structure, we find a heterotetrameric architecture composed of two RrLuc luciferases, which dock at opposite sides of the head-to-tail dimer of RrGFP. Previous studies showed that the homodimerization of fluorescent proteins brings synergistic effects, with the dimer being brighter than the sum of the monomers^25^. This could explain why the naturally evolved *Renilla* BRET complex is built on the RrGFP dimer core. The intermolecular distance between donor (CEI oxyluciferin) and acceptor (CRO fluorophore) chromophores in the BRET complex is below 3 nm edge-to-edge (3.5 nm centroid-to-centroid), favouring the Förster-type radiationless energy transfer. Furthermore, our results oppose previous beliefs that RrGFP is only a passive antenna and show that the effect of RrLuc□ RrGFP complexation on bioluminescence output is much wider. Our experiments demonstrate that the RrGFP serves not only as the colour-switchable transmitter, but it also controls the dynamical behaviour of RrLuc luciferase. HDX-MS measurements complement the crystallographic structures by demonstrating that RrGFP binding alters the conformational behaviour of RrLuc in solution. The predominant decrease in deuterium uptake within interface-proximal and cap-domain regions is consistent with protection upon complex formation and/or stabilization of more strongly hydrogen-bonded conformations. In contrast, the increased uptake observed within the A164–F181 segment indicates a localized redistribution of conformational dynamics rather than uniform rigidification of RrLuc. Because HDX-MS reflects the combined effects of hydrogen bonding, solvent accessibility, and conformational dynamics, these changes cannot be attributed to a single structural mechanism on the basis of HDX-MS alone^26^. The affected cap-domain elements have previously been implicated in active-site accessibility, substrate recognition, catalytic activity, and product release in *Renilla*-type luciferases^3,19^. Together with the molecular simulations and kinetic measurements, the HDX-MS data support a model in which RrGFP shifts the RrLuc conformational ensemble towards catalytically productive states while reducing the population of off-pathway conformations.

Although the two RrLuc-binding interfaces found in the BRET complex are structurally identical, the binding events are not. After binding the first RrLuc molecule, the second RrLuc binding event exhibits 10-fold higher binding affinity than the first one, with *K*_d_ values of ∼60 nM and ∼600 nM, respectively. In the RrLuc-bound complex, conformational changes in RrGFP were revealed. The allosteric signalling driving these structural transitions revealed here can explain the positive cooperative behaviour for binding of the second RrLuc molecule during the heterotetrameric BRET complex assembly, which has long been a neglected mystery in the past. Future work should also examine the influence of the CTZ-binding protein (RrCBP) that is supposed to be a part of the *Renilla* BRET supercomplex, in addition to potential additional regulatory functions.

Three principal parameters of the radiationless BRET efficiency are the separation, mutual orientation, and spectral overlap between the donor-acceptor pair. The co-crystal structures of the CEI-loaded RrLuc□ RrGFP complex determined here provide detailed spatial information about intermolecular dipole-dipole coupling, involving arrangement, distance, and orientation of chromophores in their proteinaceous neighbourhoods. Our results thus open exciting prospects for rational engineering of nature-inspired artificial energy-transfer and optical processes.

## Data availability

Atomic coordinates and structural factors have been deposited in the Protein Data Bank (https://www.rcsb.org/)^27^ under PDB ID accession codes: 8RZZ, 8S0G, 8S1L and 8S1V. MD simulations data were deposited in Zenodo repository with the DOI identifier: 10.5281/zenodo.22024926. HDX-MS data have been deposited in the ProteomeXchange (PX) Consortium^28^ via the Proteomics Identifications (PRIDE)^29^ partner repository with project accession: PXD080260; Token: CvPnFWqfGmqc.

## Competing interests

The authors declare no competing interests.

## Author contributions

M.M., Z.P., L.H., J.D., D.P. and M.T. conceived the study and designed the experiments. M.S., V.N. and D.P. cloned the genes and prepared the protein samples. D.P., M.T., T.B., A.D., S.S. and Z.P. performed kinetic studies. A.S., M.S., V.N., M.N., T.G. and M.M. performed protein crystallization, X-ray data collection, and structure determinations. Y.L.J. synthesized aza-luciferin analogue. N.V. and L.H. performed HDX-MS experiments. M.V. and D.P. calculated FRET efficiency. T.D.M., J.H. and D.B. performed molecular dynamics simulations. D.P., M.T., N.V., T.G. and M.M. wrote and edited the manuscript with input from all other authors.

## Funding

The study was funded by the Czech Science Foundation, project no. GX25-17329X, and the support of the RECETOX Research Infrastructure (No. LM2023069) and CZECRIN (No. LM2023049), funded by the Ministry of Education, Youth and Sports (MEYS) of the Czech Republic. This project was also supported by the European Union’s Horizon 2020 Research and Innovation Programme under grant agreement No. 857560 (CETOCOEN Excellence) and has received funding from the Horizon Europe programme under grant agreement No. 101136607 (CLARA). Additional funding was provided through the ESIF-MEYS Johannes Amos Comenius Programme under the CLARA project (No. CZ.02.01.01/00/23_029/0008437), co-financed by the European Union and MEYS. Views and opinions expressed are however those of the author(s) only and do not necessarily reflect those of the European Union or REA. Neither the European Union nor the granting authority can be held responsible for any use that may be made of the information it contains. D.P. and T.D.M. are Brno Ph.D. Talent Scholarship holders established by the South Moravian Centre for International Mobility and funded by Brno City Municipality. We acknowledge CF BIC of CIISB, Instruct-CZ Centre, supported by MEYS CR (LM2023042) and European Regional Development Fund-Project „Innovation of Czech Infrastructure for Integrative Structural Biology“ (No. CZ.02.01.01/00/23_015/0008175).

## Acknowledgments

The authors are thankful to Swiss Light Source (SLS) synchrotron members for using their beamline facilities and help during data collection. We thank Dr. Christophe Romier (IGBMC, Illkirch) for the generous gift of the pnCS plasmid.

## Methods

### Molecular cloning and mutagenesis

The genes encoding for full-length *Renilla reniformis* green fluorescent protein (RrGFP)^1^ and stabilised luciferase (RLuc8)^2^, hereafter referred to as RrLuc for clarity, were amplified by polymerase chain reaction (PCR) and cloned into bacterial co-expression vectors^3,4^ between NdeI and BamHI restriction sites. Mutants were made by megaprimer-type PCR mutagenesis and cloned into the same vectors^5^. Specifically, the genes were inserted into a pnCS vector not coding for any fusion tag, and, in parallel, the same genes were inserted into a pET21b vector in frame with a 3′-sequence coding for a C-terminal hexahistidine tag and a 3C protease cleavage site.

### Protein expression and purification

The expression plasmids were used to transform or co-transform (for co-expression assays) chemo-competent *E. coli* BL21(DE3) by a heat-shock method. Several colonies were transferred into 10 mL of Luria-Bertani (LB) medium supplemented with ampicillin (100 µg/mL), or with both ampicillin (100 µg/mL) and spectinomycin (100 µg/mL), and cultivated for 4-5 hours at 37°C and 200 rpm. The preculture was then added into 2 L of 2×LB medium (100 µg/mL of ampicillin) and cultivated at 37°C and 200 rpm until OD_600_ ≥ 1.5. After that, expression was induced by adding isopropyl β-D-1-thiogalactopyranoside (IPTG) to the final concentration of 200 µM. Then, cells were incubated at 20°C and 150 rpm for approximately 16 hours, and harvested by centrifugation (3,220 × g, 4°C, 25 min). Pellets from three litres of culture were resuspended in ∼40 mL of buffer (5 mM KCl, 10 mM Tris, pH = 7.5) and stored at -70°C.

Cell mass was thawed and mixed with 100 µL of DNase (1 mg/mL) and sonicated using Sonic Dismembrator Model 705 (Fisher Scientific, USA) in 4 × 2 min cycles (5 s pulse, 5 s pause) with 50% amplitude. The sonicated lysate was diluted with ∼80 mL of buffer (5 mM KCl, 10 mM Tris, pH = 7.5) and clarified by centrifugation (19,500 × g, 4°C, 1 hour). Supernatants containing the recombinant proteins were purified using Talon metal affinity resin (Clontech). To release the His-tagged proteins and complexes from the resin, the samples were treated with 3C protease overnight at 4°C. The next day, the eluted proteins were separated by gel filtration chromatography on Äkta FPLC (Cytiva) equipped with HiLoad^™^ 16/600 Superdex^™^ 200 pg column (Cytiva) equilibrated with buffer (5 mM KCl, 10 mM Tris-HCl buffer, pH 7.5). Purified proteins were concentrated to final concentrations using Centrifugal Filter Units Amicon^®^ Ultra-15 Ultracel^®^-10K (Merck Millipore). Protein purity was verified by SDS-PAGE. Concentration of protein samples was measured using DeNovix^®^ DS-11 Spectrophotometer (DeNovix).

### Spectroluminometric analysis

All emission spectra were obtained at 25°C, using a spectrofluorometer FluoroMax Plus (HORIBA, Japan), in 5 mM KCl, 10 mM Tris-HCl buffer (pH 7.50). This buffer was chosen as an environment with ionic strength (∼11.5 mM) sufficiently low to be suitable for the interaction between RrLuc and RrGFP^6^, yet high enough not to interfere with other methods used.

RrGFP fluorescence emission spectra were obtained for 10.0 µM RrGFP in 5 mM KCl, 10 mM Tris-HCl (pH 7.50) buffer in a quartz cuvette. RrGFP sample was excited at 470 nm, and fluorescence emission spectra were collected in tetraplicate over the range of 370–680 nm with a 2 nm interval. Luminescence emission spectra were measured for 0.4 µM RrLuc in 5 mM KCl, 10 mM Tris-HCl buffer (pH 7.50) in a quartz cuvette either alone or mixed with RrGFP in 0.125-, 0.25-, 0.5-, 1-, 2-, 4-, 8-, and 16-times molar ratio to RrLuc. The luminescence reaction was started by the addition of 20 µL of ∼500 µM ethanolic stock solution of coelenterazine (CTZ). Emission spectra of RrLuc luminescence in the 16-fold molar excess of RrGFP were also determined in the presence of 300 and 650 mM NaCl. Luminescence spectra were measured in triplicate over the range of 370–680 nm with a 2 nm interval.

The dependence of the ratio of emission intensity at 470 and 509 nm on RrGFP concentration, a measure of the interaction between RrLuc and RrGFP, was fitted with a model of non-cooperative binding (**Eq. 1**) by software GraphPad Prism (GraphPad Software, USA):

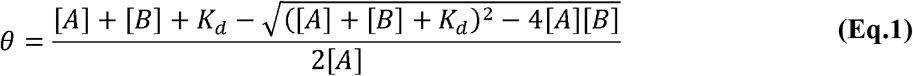

where [A] represents the total concentration of observed interaction partner A of a constant concentration, [B] is the total concentration of interaction partner B of variable concentration, and θ represents the partial amount of the interaction partner A, which participates in the interaction with a dissociation constant *K*_d_ at a given concentration of both interaction partners.

### MST-TRIC measurements

A 16-point 5·0.6^0–15^ μM dilution series of RrLuc was prepared in a mixture with 10 nM RrGFP. Samples were loaded into glass capillaries and analysed by Monolith NT.115 (NanoTemper, Germany). The fluorescent profiles of the capillaries tempered to 25°C were determined to assess the adhesion of RrGFP to the capillary walls. Capillaries were subsequently locally heated by an IR laser (λ = 1475 ± 15 nm, P = 120 mW) and monitored for a change in RrGFP fluorescence (λ_ex_ = 465–490 nm, λ_em_ = 500–550 nm) in the heated location due to the MST-TRIC (Microscale thermophoresis and temperature related intensity change) effect. The same measurement was repeated in the presence of a 12.5-fold molar excess of CTZ and substrate analogue azaCTZ relative to the concentration of RrLuc. The aza-luciferin analogue (AzaCTZ) was synthesized as described previously^5^. All measurements were conducted in 5 mM KCl, 10 mM Tris-HCl buffer (pH 7.5) in triplicate.

Affinity analyses (NanoTemper, Germany) were performed using MST-TRIC data collected between 1.5 and 2.5 s after the start of the measurement. The measured dependencies of RrGFP fluorescence on RrLuc concentration were first fit to a non-cooperative binding model (**Eq. 1**), expecting independent binding of RrLuc to the two binding sites of RrGFP. Later, models considering cooperativity were found to be a better option, namely, the Hill-Langmuir model (**Eq. 2**) and the Adair-Klotz model (**Eq. 3-4**).

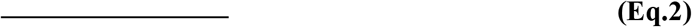

The Hill-Langmuir model assumes cooperativity in the simultaneous binding of multiple molecules of interaction partner B to a single molecule of interaction partner A, as reflected in the Hill coefficient *n*_H_. Symbols θ and [B] represent the same quantities as in **Eq. 1**, and *K*_d,app_ is the apparent dissociation constant of the interaction, which is the value of [B] causing half-maximal occupation of binding sites of the interaction partner A^2^.

Another model used for data analysis expects the sequential binding of two molecules of interaction partner B onto one molecule of interaction partner A according to the scheme (**Eq. 3**):

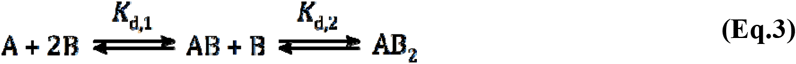

which is described by **Eq. 4** derived from the Adair-Klotz model:

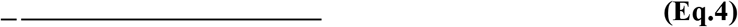

where symbols θ and [B] represent the same quantities as in **Eqs. 1** and **2** and *K*_d,1_ and *K*_d,2_ are the macroscopic dissociation constants of the sequential binding steps. The value of *K*_d,app_ is determined as the geometric average of the determined dissociation constants.

### Temperature-dependent transient kinetics of CTZ binding

Transient kinetic traces of CTZ binding to free RrLuc luciferase or the RrLuc−RrGFP complex were collected at temperatures ranging from 8 to 21°C after rapidly mixing the components using an advanced continuous-flow microfluidic chip. The method allows high-throughput data collection with minimal sample consumption and rapid heat-transfer in a setup described and published previously^7^. The measurement was initiated by mixing the RrLuc or RrLuc−RrGFP solution with the CTZ solution at a total flow rate of 2.1 µL.min^-1^ and then monitored for the change of CTZ fluorescence upon binding using a 488 nm long-pass emission filter after excitation at 405 nm. The experiment was performed in a 10 mM Tris-HCl, 5 mM KCl buffer (pH 7.5). For measurements with free RrLuc, the final concentration of CTZ after mixing was kept constant at 27 µM while the luciferase concentration varied from 8 to 71 µM. For measurements with RrLuc−RrGFP, the final concentration of the complex after mixing was kept constant at 1.25 µM while the concentration of CTZ varied from 1.7 to 15 µM.

### Stopped-flow transient kinetics of CTZ binding

Transient kinetic traces of CTZ binding to either free RrLuc or the RrLuc−RrGFP complex were additionally obtained in a conventional stopped-flow experiment setup, using the instrument Stopped-Flow SFM-3000 equipped with a rapid kinetics fluorescence spectrometer MOS-200 and a PMT detector (BioLogic, France). The experiment was performed in a 10 mM Tris-HCl, 5 mM KCl buffer (pH 7.5), at 15°C. The measurement was initiated by rapid mixing of RrLuc (final concentration 0.5 µM) or a mixture of RrLuc and RrGFP (final concentrations 0.5 µM and 5 µM, respectively) with a 2:1 dilution series of CTZ (final concentrations ranging from 0.8 µM to 30 µM) in a 1:1 volume ratio (75 + 75 µl) at a total flow rate of 11.50 mL s^−1^ and deadtime of 0.3 ms (µFC−08 cuvette 0.8 mm, valve lead time 3.3 ms), including a control experiment of rapid-mixing RrLuc or RrLuc−RrGFP with assay buffer without any CTZ content. After the mixing, the CTZ binding was monitored through a change in the RrLuc tryptophan fluorescence (excitation: 280 nm, emission: 340 ± 13 nm band-pass filter) for 0.3 s in 0.5 ms intervals. Each data trace was obtained as an average of 10-15 subsequent identical rapid-mixing experiments.

### Data analysis for pre-steady-state kinetics of CTZ binding

Global numerical analysis of kinetic data was performed using KinTek Explorer 10 (KinTek Corporation, USA)^8–10^. The software allows the input of a given kinetic model via a simple text description, and the program automatically derives the differential equations needed for numerical integration. The induced-fit mechanism (**Eq. 5**), identified for the RrLuc enzyme^11^, was used as the input kinetic model:

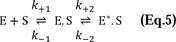

In the case of the on-chip measurement using CTZ fluorescence, the previously observed dependence of increasing CTZ fluorescence intensity over time upon binding into free RrLuc was extended with two more parameters due to the presence of RrGFP: (*i*) background fluorescence of RrGFP itself in the region of measurement, and (*ii*) fluorescence decrease at later kinetic stages due to the fluorescence resonance energy transfer (FRET) shifting emission of the E*.S complex out of the measurement region. As a result, the fluorescence changes at temperature *T* were described with **Eq. 6**:

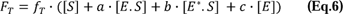

In the case of the conventional stopped-flow experiments using the RrLuc tryptophan fluorescence, no such mechanism extensions were needed. RrGFP presence, when applicable, introduced only a small constant measurement background increase due to the lack of tryptophan residues in the RrGFP sequence.

Numerical integration of rate equations searching a set of kinetic parameters that produce a minimum χ^2^ value was performed using the Bulirsch–Stoer algorithm with adaptive step size, and nonlinear regression to fit data was based on the Levenberg–Marquardt method. Residuals were normalized by sigma value for each data point. The standard error (S.E.) was calculated from the covariance matrix during nonlinear regression. To account for fluctuations in experimental data, substrate concentrations were allowed to be slightly adjusted (±10%) to obtain the best fits. The final values of the elementary rate constants (*k*_+1_, *k*_-1_, *k*_+2_, and *k*_-2_) were calculated by fitting all the raw kinetic data globally (separately for the on-chip and conventional stopped-flow experiments). For on-chip measurements in variable temperature, activation energies (*E*_a,+1_, *E*_a,-1_, *E*_a,+2_, and *E*_a,-2_) were evaluated in the same fashion.

In addition to standard error values, a more rigorous analysis of the variation of the determined parameters was accomplished by confidence contour analysis using FitSpace Explorer (KinTek Corporation, USA). In these analyses, the lower and upper limits for each parameter were derived from the confidence contour obtained after setting χ^2^ threshold at 0.95.

The values of the apparent dissociation constant (*K*_d,ES(app)_) were calculated according to **Eq. 7**.

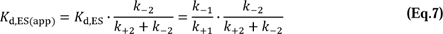

Fluorescence signal changes upon mixing RrLuc–RrGFP with the CTZ substrate exhibited two kinetic phases in accordance with previous observations, corresponding to the induced-fit binding mechanism^6^. However, the second kinetic phase of RrLuc–RrGFP kinetics yielded a decreasing trend in time, as opposed to solely increasing substrate fluorescence observed upon binding by free luciferase. The drop of the fluorescence was assigned to the fluorescence resonance energy transfer (FRET) from the substrate to RrGFP whose absorbance overlaps with the substrate emission region used for kinetic experiments. We anticipate that the substrate fluorescence initially increases due to the binding to a more hydrophobic and spatially restricted active site (observed also for free luciferase). However, upon a follow-up conformational change (second step of the binding mechanism), the substrate gets into the vicinity of RrGFP. At this point, RrGFP starts absorbing the substrate fluorescence, so instead of detecting additional fluorescence increase, typical for free RrLuc, the signal starts decreasing because the emitted light is red-shifted out of the measurement region due to the discussed FRET. When the fluorescence decrease caused by FRET upon the conformational change was assumed in the kinetic model (**Eq. 6**), trends corresponding to the induced-fit binding mechanism could be obtained.

### Steady-state kinetics of CTZ conversion

Steady-state kinetic parameters of RrLuc catalysis were determined for free RrLuc and RrLuc complexed with RrGFP using a microplate reader FLUOstar Omega (BMG Labtech). All reactions were performed in white microtiter plates, in a 5 mM KCl, 10 mM Tris-HCl buffer (pH 7.5). CTZ solutions (4.4, 2.2, 1.1, 0.55, and 0.28 μM) were prepared immediately before the measurements by diluting an ethanolic stock solution of CTZ into 3 mL of the reaction buffer. The final concentration range of CTZ (0.25–3.96 μM) was chosen around the expected value of the Michaelis constant *K*_m_. The concentration of RrLuc (0.02 μM) was chosen as a compromise between a requirement for the enzyme concentration to be low to ensure that every molecule of RrLuc catalyses the conversion of at least 10 molecules of CTZ on average, even with the lowest CTZ concentration, and a requirement for the concentration to be high enough so that the total conversion of CTZ proceeds in a reasonable timeframe, making the amount of CTZ that autooxidises negligible when compared to the amount of CTZ oxidised by RrLuc. The luminescent reaction was initiated by an automatic injection of 225 μL of CTZ solution into a well with 25 μL of 0.2 μM RrLuc or 0.2 μM RrLuc + 36.0 μM RrGFP; the concentration of RrGFP was chosen significantly above the determined RrLuc−RrGFP interaction *K*_d_ to ensure the complete complexation of RrLuc with RrGFP. A broad spectrum of luminescence (240–740 nm) of the reaction mixture was monitored by a CCD detector in 1s intervals with 1s exposure time until the luminescent signal fell below 0.5 % of its maximal measured value. The described measurement was performed in triplicate for all prepared CTZ solutions. The luminescent background of the reactions was determined by the same method for 225 μL of CTZ solution injected into a 25 μL of the reaction buffer and also for 225 μL of reaction buffer injected into 25 μL of RrLuc or RrLuc−RrGFP solution. All measurements were performed at 37°C with a constant gain setting of the detector.

### Data analysis for steady-state kinetics of CTZ conversion

The measured luminescence signal was transformed into a dependency of total luminescence intensity (TLI) on the reaction time, which is directly proportional to the dependency of the amount of product created on the reaction time. The transformed data were evaluated numerically using KinTek Explorer (KinTek Corporation, USA)^9^, and the reliability of the obtained parameter values was determined by confidence contour analysis using FitSpace Explorer (KinTek Corporation, USA)^9^, as described above for pre-steady-state kinetics of CTZ binding. (**Extended Data Fig. 5**).

Kinetic parameters of the RrLuc catalytic reaction for free RrLuc and the complex RrLuc–RrGFP were determined according to the reaction model (**Eq. 8**):

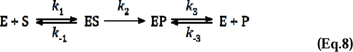

where E, S, and P represent free enzyme, substrate, and product, respectively. ES and EP stand for complexes of enzyme-substrate and enzyme-product, respectively, and *k*_n_ represents the rate constants of individual reaction steps. Analysis was performed under the assumption of the irreversibility of the luciferase reaction (*k*_-2_ = 0 s^-1^) and under rapid equilibrium of formation of enzyme-substrate and enzyme-product complexes limited only by diffusion (*k*_1_ = *k*_-3_ = 100 μM^-1^ s^-1^). The interaction of RrLuc and RrGFP was not factored into the reaction scheme, considering the determined nonsignificant effect of azaCTZ or CTZ presence on the interaction between RrLuc and RrGFP. Since the mixing of RrLuc and RrGFP solutions took place before the reaction itself, it is therefore assumed that at the moment of reaction start, all RrLuc is already complexed with RrGFP and no additional binding event needs to happen for the complex to be able of catalysis. Under these assumptions, ratio *k*_-1_/*k*_1_ represents the Michaelis constant (*K*_m_), ratio *k*_3_/*k*_-3_ equals the dissociation constant of enzyme-product complex (*K*_d,EP_) expressing the strength of product inhibition, and *k*_2_ equals the turnover number of the reaction (*k*_cat_). Catalytic efficiency *k*_cat_/*K*_m_ was determined as a fitted parameter by assigning a value of 0 s^-1^ to *k*_-1_, which causes parameter *k*_1_ to equal *k*_cat_/*K*_m_^9^

Apart from kinetic constants, one of the fitted parameters was the relative specific total luminescence intensity (TLI_s,rel_) as a factor of proportionality between the transformed luminescence signal and reaction time dependency of the formed product. Evaluation of this parameter was made possible by monitoring the complete conversion of the substrate. This parameter, considering the type of detector used, is directly proportional to the total energy of emitted photons, and, therefore, representative of the relative quantum yield of the reaction. Due to the reaction setup, TLI_s,rel_ is not as reliably determinable as the evaluated catalytic parameters, nor is it comparable to previously published data on the effect of RrGFP on the quantum yield of RrLuc bioluminescence. However, it represents a good estimate of relative differences in quantum yields across systems analysed by steady-state kinetics within this study.

### Protein crystallizations

All crystallisation experiments were done at 20°C using the hanging-drop vapour-diffusion method in EasyXtal 15-well plates (Qiagen) with drops equilibrated against 500 uL of reservoir solution. For coelenteramide (CEI) bound RrLuc−RrGFP complex (RrLuc/CEI−RrGFP), the RrLuc−RrGFP reconstituted via co-expression in *E*. *coli* was concentrated to 30 mg/mL and activated by adding substrate CTZ in 1:4 molar protein-ligand ratio. After incubation for 30 minutes, the complex was subjected to crystallisation experiments. Diffraction-grade crystals were obtained after mixing 1.5 μL of RrLuc/CEI−RrGFP complex with 1 μL of the precipitant solution consisting of 0.2 M sodium tartrate dibasic hydrate and 20% (w/v) PEG 3350, or 0.1 M MES (pH 6.5) and 15% (w/v) PEG 4000, or 7% (w/v) T-mate (pH 7), 0.1 M Bicine (pH 9), 22.5% (v/v) PEG Smear High and 10% (v/v) ethylene glycol (Molecular Dimensions).

For apo-RrGFP crystals, the RrGFP protein was concentrated to ∼15 mg/mL. 1 μL of protein was mixed with the precipitant solution consisting of 0.2 M MgCl_2_, 0.1 M Hepes pH 6.5 and 20% PEG 3350. All crystals were fished out, cryo-protected in the corresponding reservoir solutions supplemented with 20% glycerol, and flash-frozen in liquid nitrogen for X-ray data collection.

### X-ray data collection and structure determinations

Diffraction data were collected at Swiss Light Source beamline X06DA at a wavelength of 1.0 Å, using a Pilatus 2M-F detector. The data were processed using XDS^12^ and Aimless^13^. Structures were solved by molecular replacement using Phaser^14^ in the Phenix^15^ software suite. Previously determined structures of RrLuc (PDB ID: 6YN2^11^) and RrGFP (PDB ID: 2RH7^1^) were used as search models. Iterative refinements were done using phenix.refine in Phenix^15^. Manual model building and refinement were performed in Coot^16^. The refinement statistics are listed in **Extended Data Table 1**. The structures were visualized by PyMOL^17^, using DSSP algorithm^18^ for the classification of the protein secondary structures. The PDBePISA server^19^ was used to identify the interactions at the dimerisation interface and interaction interface between RrLuc and RrGFP, as well as their corresponding distances, calculate the interface area, and solvation free energy gain upon formation of the interface (Δ^i^G).

### Calculation of FRET efficiency

FRET efficiency was calculated for five RrLuc-bound CEI oxyluciferin molecules (FRET donors) and all six RrGFP CRO fluorophores (FRET acceptors) captured within the herein solved RrLuc−RrGFP complex structure (PDB ID: 8RZZ). For each meaningful donor-acceptor pair, we extracted the distance *r* as the centroid-to-centroid distance between the CEI and CRO moieties, angle θ_T_ between the donor and acceptor transition dipole moments, angle θ_D_ between the donor transition dipole moment and the donor-acceptor intermolecular axis, and angle θ_A_ between the acceptor transition dipole moment and the donor-acceptor intermolecular axis. The orientations of transition dipole moments of the donor and acceptor with respect to their molecular structures were obtained from literature^20^.

To assess the likelihood of FRET in particular donor-acceptor pairs, we used the Förster resonant energy transfer (FRET) model^21^, assuming that the spectral properties (frequency-dependent extinction coefficients) of the acceptors are comparable, and the difference in the FRET rates between the pathways is thus determined only by the orientation factor κ^2^ and the donor-acceptor distance *r*. Using this assumption and the standard formula for the FRET rate, we define a FRET efficiency factor *f*_DA_ as *f*_DA_ = κ^2^/*r*^6^; the value of κ was calculated as κ = cos(θ_T_) − 3 cos(θ_D_) cosθ_A_. The acceptor preference is calculated as a ratio of FRET efficiency factor *f*_DA_ for the particular CRO acceptor and the other CRO acceptor in the same RrGFP dimer. The acceptor preference can be decomposed into the orientation advantage, equal to the ratio of the FRET orientation factors κ^2^ of the proximal and distal CRO acceptors (κ^2^_proximal_/κ^2^_distal_), and the distance advantage, calculated as the ratio of the sixth powers of the distal and proximal acceptor distances *r* (*r*^6^_distal_/r^6^_proximal_); the acceptor preference is the product of the orientation and distance advantages. The acceptor preference, orientation advantage, and distance advantage are defined relative to the proximal acceptor; for the distal acceptor, the corresponding values are the reciprocals of the defined ratios. The orientation enhancement is the fold-improvement in *f*_DA_ for a particular CRO acceptor over an imaginary equidistant acceptor with random orientation in space (κ^2^ = ⅔).

### Hydrogen-deuterium exchange mass spectrometry

HDX-MS analysis of the RrLuc luciferase was conducted in both unbound and RrGFP-bound forms, using the procedures described previously^22,23^. Initially, undeuterated sample preparation for RrLuc was optimised and analysed independently using a mass spectrometer operated in data-dependent MS/MS mode to identify peptides. These peptides provided essential information regarding their abundance and sequence coverage. Subsequently, the optimised sample preparation and analytical conditions were applied to HDX-MS analyses. Control undeuterated samples of RrLuc, both unbound and RrGFP-bound, were diluted to a final concentration of 1 µM in a basic H_2_O buffer (10 mM Tris–base and 5 mM KCl, pH 7.5). For deuterated samples, a buffer of identical composition (pD 7.5, pH 7.1) was prepared. Each HDX time point was prepared by mixing the RrLuc and RrGFP at a 1:50 molar ratio. The mixture was pre-incubated for 30 minutes at room temperature and then diluted with deuterated buffer to achieve a final composition of 50% D_2_O. Four distinct time points (60 s, 300 s, 600 s, and 1800 s) were collected by quenching the reaction by adding an ice-cold quenching buffer (4 M urea, 0.5M TCEP, 1 M glycine, and 1 mg/mL pepsin, pH 2.3) in a 1:1 ratio, followed by a 3-minute incubation at room temperature and rapid freezing in liquid nitrogen.

Each sample was thawed and injected into an LC system (UltiMate 3000 RSLCnano, Thermo Scientific Dionex, Massachusetts, USA) equipped with an immobilised pepsin and nepenthesin enzymatic column (15 µL bed volume, flow rate of 100 µL/min, 2% acetonitrile/0.05% trifluoroacetic acid). Peptides were trapped and desalted online using a peptide microtrap (Michrom Bioresources, Auburn, CA, USA) for 3 minutes at a flow rate of 100 µL/min. Subsequently, the peptides were eluted onto an analytical column (Jupiter C18, 1.0 × 50 mm, 5 µm, 300 Å, Phenomenex, CA, USA) and separated using a 29-minute linear gradient elution from 10% to 40% of mobile phase B (80% acetonitrile/0.08% formic acid) in mobile phase A (0.1% formic acid in water). The enzymatic column with immobilised pepsin and nepenthesin, trap cartridge, and analytical column were maintained at 1°C.

### Mass spectrometry and data analysis

Mass spectrometric analysis was performed using an Orbitrap Elite mass spectrometer (Thermo Fisher Scientific, Massachusetts, USA) with ESI ionisation, integrated with a robotic system based on the HTS-XT platform (CTC Analytics, Zwingen, Switzerland). The instrument operated in data-dependent mode for peptide mapping (HPLC–MS/MS). Each MS scan was followed by MS/MS scans of the three most intense ions from the CID and HCD fragmentation spectra. Tandem mass spectra were searched using SequestHT against the cRAP protein database (ftp://ftp.thegpm.org/fasta/cRAP), which included sequences of RrLuc and RrGFP. Search settings were as follows: precursor ion mass tolerance of 10 ppm, fragment ion mass tolerance of 0.6 Da, no enzyme specificity, and no fixed or variable modifications. The peptide identification false discovery rate was set to 1%. Sequence coverage was analysed using Proteome Discoverer software version 1.4 (Thermo Fisher Scientific, Massachusetts, USA). Deuterated sample analysis was conducted in LC–MS mode with ion detection in the orbital ion trap. MS raw files, along with the identified peptide pool (including amino acid sequences, retention times, XCorr values, and ion charges), were processed using HDExaminer version 2.2 (Sierra Analytics, Modesto, CA, USA). The software assessed peptide deuteration kinetics and generated uptake plots displaying peptide deuteration over time, with confidence levels categorized as high or medium (accepted) or low (rejected). Relative fractional uptake (RFU) values for individual amino acids were calculated using PyHDX version 0.4.3^24^. For both unbound and RrGFP-bound RrLuc, RFU values were determined by weighted averaging of overlapping peptides, with weighting based on the inverse peptide length. Results were visualised as RFU plot and differential RFU mapped on the structure and sequence. Evaluated data, peptide-level uptake and RFU/ΔRFU values, are provided in **Supplementary Source Data** file. Raw HDX-MS data are deposited in the PRIDE repository, with project accession: PXD080260; Token: CvPnFWqfGmqc.

### Ligand preparation for computational studies

The structure of CEI was prepared using Avogadro 1.2.0 software^25^: the multiplicity of the bonds was edited to match the keto form, all missing hydrogens were added, and the structure was minimized by the steepest descent algorithm in the Auto Optimize tool of Avogadro, using the Universal Force Field (UFF). Next, the *antechamber* module of AmberTools16^26^ was used to calculate the charges for the ligand, add the atom types of the Amber force field, and compile them in a PREPI parameters file. Also, the *parmchk2* tool from AmberTools16 was used to create an additional FRCMOD parameter file to compensate for any missing parameters. The structure of the RrGFP chromophore (CRO) and the corresponding PREPI and FRCMOD parameter files were prepared following the Amber tutorial.

### Adaptive sampling molecular dynamics analysis

Release of CEI from the RrLuc–RrGFP complex was simulated using adaptive sampling from the High Throughput Molecular Dynamics (HTMD)^24^ package. System preparation was carried out as previously described, with minor changes^27^. Briefly, the system was protonated with PROPKA 2.0^28^ at pH 7.5. Minimization and equilibration were extended to 10 ns of equilibration at a temperature of 298K with a scaling factor of 0.8333 for the 1-4 electrostatic interactions. A 9 Å cut-off was employed for long-range interactions using the particle mesh Ewald method. 40 epochs of 10 parallel MD simulations were performed, each 50 ns in length, for a total of 20 µs. For product release, adaptive sampling was performed using the distance of the N1 atom of CEI and the □-carbon (CG) atom of catalytic residue D120 as the adaptive metric, together with time-lagged independent component analysis (tICA)^29^ projection in 1 dimension. Unsuccessful simulations shorter than 50 ns were omitted, after which water and ion molecules were filtered out using AmberTools16. Markov state model (MSM) construction and kinetic analysis were subsequently performed as previously described with a lag time of 17 ns^30^.

### Accession codes

X-ray crystallographic coordinates and structure factor files have been deposited in Protein Data Bank^31^ under accession numbers: 8RZZ, 8S0G, 8S1L and 8S1V. HDX-MS data have been deposited in the ProteomeXchange (PX) Consortium^32^ via the Proteomics Identifications (PRIDE)^33^ partner repository with project accession: PXD080260; Token: CvPnFWqfGmqc.

## Notes

### Competing Interest Statement

The authors have declared no competing interest.

