## Extended Data for "Structural mechanism governing radiationless energy transfer in *Renilla* bioluminescence"

**Extended Data Table 1.** **Affinity binding parameters retrieved from thermophoretic measurements using both non-cooperative and cooperative models.** *K*_d_ – dissociation constant, *K*_d,app_ – apparent dissociation constant, *n*_H_ – Hill coefficient, *K*_d,1_ and *K*_d,2_ – macroscopic dissociation constants of the first and second sequential binding event, respectively, R^2^ – coefficient of determination expressing the goodness of the fit. *K*_d,1_/*K*_d,2_ ratio obtained from Adair-Klotz (~2.5), exceeds the value typical for non-cooperative binding (0.25) 10-fold. Macroscopic dissociation constants are influenced by statistical factors: the first binding of RrLuc to RrGFP is 2-fold more probable than the second binding, while the release of RrLuc from the full 2:2 complex is 2-fold more likely than from the partial 1:2 complex. Values of microscopic dissociation constants of the first and second binding events were obtained by correction of fitted macroscopic dissociation constants *K*_d,1_ and *K*_d,2_ as 604 ± 54 nM and 62 ± 6 nM, respectively.

| Fitting model | Non-cooperative binding model | | Hill-Langmuir model | | | Adair-Klotz model | | | | |
| --- | --- | --- | --- | --- | --- | --- | --- | --- | --- | --- |
| Sample composition | *K*_d_ / nM | R^2^ | *K*_d,app_ / nM | *n*_H_ | R^2^ | *K*_d,1_ / nM | *K*_d,2_ / nM | *K*_d,1_/ *K*_d,2_ | *K*_d,app_ / nM | R^2^ |
| RrLuc−RrGFP | 194 ± 31 | 0.985 | 191 ± 15 | 1.51 ± 0.16 | 0.994 | 302 ± 27 | 123 ± 12 | 2.5 ± 0.3 | 192 ± 18 | 0.994 |
| RrLuc−RrGFP + azaCTZ | 206 ± 51 | 0.967 | 183 ± 30 | 1.55 ± 0.34 | 0.987 | 331 ± 59 | 103 ± 20 | 3.2 ± 0.6 | 184 ± 34 | 0.987 |
| RrLuc−RrGFP + CTZ | 201 ± 37 | 0.981 | 181 ± 18 | 1.56 ± 0.21 | 0.990 | 310 ± 45 | 106 ± 13 | 2.9 ± 0.4 | 181 ± 24 | 0.989 |

**Extended Data Table 2. Crystallographic data collection and refinement statistics.**

|  | RrLuc8‒RrGFP (pH 9) | RrLuc8‒RrGFP | RrLuc8‒RrGFP (pH 6) | RrGFP |
| --- | --- | --- | --- | --- |
| Data collection |  |  |  |  |
| Wavelength (Å) | 1 | 1 | 1 | 1 |
| Space group | *P*32 | *P*32 | *P*32 | *P*4_3_2_1_2 |
| Cell dimensions |  |  |  |  |
| a, b, c (Å) | 97.99, 97.99, 361.99 | 96.88, 96.88, 361.46 | 95.93, 95.93, 357.13 | 82.43, 82.43, 82.22 |
| α, β, γ (°) | 90, 90, 120 | 90, 90, 120 | 90, 90, 120 | 90, 90, 90 |
| Resolution (Å) | 49.17 - 2.30 (2.38 - 2.30) | 48.43 - 2.38 (2.46 – 2.38) | 46.32 - 2.4 (2.48 - 2.4) | 36.84 - 1.94 (2.01 - 1.94) |
| Total reflections | 1,798,891 (175,467) | 612,940 (28,036) | 1,514,543 (142,294) | 232,792 (8,151) |
| Unique reflections | 172,769 (17,230) | 152,020 (15,053) | 143,749 (14,312) | 21,390 (2,034) |
| Rmerge | 0.177 (1.942) | 0.094 (1.116) | 0.222 (4.653) | 0.039 (0.882) |
| I/σI | 12.7 (1.3) | 9.0 (0.9) | 9.9 (0.6) | 29.5 (1.2) |
| Completeness (%) | 99.9 (99.6) | 99.5 (97.8) | 99.0 (96.1) | 99.5 (96.6) |
| Multiplicity | 10.4 (10.2) | 4.0 (3.8) | 10.5 (9.9) | 10.9 (4.0) |
| CC(1/2) | 0.998 (0.576) | 0.997 (0.482) | 0.998 (0.369) | 1.0 (0.654) |
| Wilson B-factor | 43.34 | 46.75 | 55.7 | 44.19 |
| Refinement |  |  |  |  |
| Resolution (Å) | 49.17 - 2.30 (2.38 - 2.30) | 48.01 - 2.38 (2.46 – 2.38) | 46.32 - 2.4 (2.48 - 2.4) | 36.84 - 1.94 (2.01 - 1.94) |
| No. reflections | 172,769 (17,217) | 152,020 (14,957) | 143,749 (13,814) | 21,390 (2,031) |
| No. reflections for R-free | 8,471 (865) | 7,219 (750) | 7,058 (631) | 1,151 (118) |
| Rwork / Rfree (%) | 22.67 / 25.65 | 21.98 / 28.13 | 26.76 / 33.95 | 20.39 / 24.92 |
| No. atoms |  |  |  |  |
| Protein | 25,998 | 25,911 | 17,471 | 1,816 |
| Ligand | 281 | 157 | 208 | 33 |
| Water | 648 | 373 | 93 | 124 |
| B-factors | 45.11 | 57.54 | 78.78 | 56.37 |
| Protein | 45.08 | 57.57 | 78.74 | 56.28 |
| Ligand | 50.05 | 61.01 | 85.70 | 59.35 |
| Water | 43.99 | 54.30 | 69.45 | 56.93 |
| R.m.s. deviations |  |  |  |  |
| Bond lengths (Å) | 0.003 | 0.004 | 0.011 | 0.002 |
| Bond angles (°) | 0.62 | 0.72 | 1.5 | 0.65 |
| Ramachandran favored (%) | 93.18 | 93.24 | 80.50 | 99.56 |
| Ramachandran allowed (%) | 5.97 | 5.57 | 13.85 | 0 |
| Ramachandran outliers (%) | 0.85 | 1.19 | 5.64 | 0.44 |
| PDB ID code | 8RZZ | 8S0G | 8S1L | 8S1V |
| Values in parentheses are for the highest-resolution shell. | | | |  |

**Extended Data Table 3. Calculated FRET parameters.** Values were determined for five RrLuc-bound CEI oxyluciferin molecules (FRET donors) and all six RrGFP CRO fluorophores (FRET acceptors) captured within the RrLuc−RrGFP complex structure (PDB ID: 8RZZ). The error shown is the S.D. of the five individually determined values. “Proximal acceptor” denotes the CRO fluorophore of the RrGFP molecule in the main-interface interaction with the RrLuc binding the CEI oxyluciferin, while “distal acceptor” is the CRO fluorophore of the other RrGFP molecule present in the 2:2 complex. *r*: centroid-to-centroid distance between the CEI donor and CRO acceptor chromophores; ***θ***_T_: angle between the donor and acceptor transition dipole moments; ***θ***_D_: angle between the donor transition dipole moment and the donor-acceptor intermolecular axis; ***θ***_A_: angle between the acceptor transition dipole moment and the donor-acceptor intermolecular axis; κ^2^: FRET orientation factor; *f*_DA_: FRET efficiency factor. Orientation enhancement is the fold-improvement in *f*_DA_ for a particular CRO acceptor over an equidistant acceptor with random orientation in space (κ^2^ = ⅔), highlighting the importance of the evolved CEI-CRO orientation to the FRET efficiency in the RrLuc-RrGFP complex. The acceptor preference is calculated as a ratio of FRET efficiency factor *f*_DA_ for the particular CRO acceptor and the other CRO acceptor in the same RrGFP dimer. The acceptor preference can be decomposed into the orientation advantage, equal to the ratio of the FRET orientation factors κ^2^ of the proximal and distal acceptors (κ^2^_proximal_/κ^2^_distal_), and the distance advantage, calculated as the ratio of the sixth powers of the distal and proximal acceptor distances *r* (*r*^6^_distal_/r^6^_proximal_); the acceptor preference is the product of the orientation and distance advantages. The acceptor preference, orientation advantage, and distance advantage are defined relative to the proximal acceptor; for the distal acceptor, the corresponding values are the reciprocals of the defined ratios.

| Parameter | Proximal acceptor | | Distal acceptor | |
| --- | --- | --- | --- | --- |
|  | Value | Spread | Value | Spread |
| *r* / nm | 3.49 ± 0.04 | 3.44-3.53 | 4.21 ± 0.04 | 4.18-4.28 |
| *θ*_T_ / rad | 1.60 ± 0.05 | 1.54-1.65 | 1.88 ± 0.05 | 1.82-1.94 |
| *θ*_D_ / rad | 2.56 ± 0.07 | 2.47-2.63 | 2.48 ± 0.04 | 2.44-2.53 |
| *θ*_A_ / rad | 2.02 ± 0.02 | 2.00-2.04 | 1.70 ± 0.02 | 1.67-1.72 |
| κ^2^ | 1.24 ± 0.04 | 1.20-1.29 | 0.37 ± 0.08 | 0.28-0.47 |
| f_DA_ (×10^4^) | 6.88 ± 0.43 | 6.55-7.61 | 0.66 ± 0.17 | 0.49-0.86 |
| Orientation enhancement (vs. random orientation) | 1.86 ± 0.06 | 1.80-1.94 | 0.55 ± 0.12 | 0.42-0.70 |
| Acceptor preference | 10.81 ± 2.23 | 8.01-13.69 | 0.10 ± 0.02 | 0.07-0.12 |
| Orientation advantage | 3.51 ± 0.87 | 2.58 - 4.63 | 0.30 ± 0.07 | 0.22 - 0.39 |
| Distance advantage | 3.11 ± 0.22 | 2.84 - 3.40 | 0.32 ± 0.02 | 0.29 - 0.35 |

**Extended Data Table 4.** **Hydrogen bonds and salt bridges between RrLuc and RrGFP. Residues forming non-polar or hydrophobic contacts of the RrLuc−RrGFP interaction interface.** Residues of RrLuc: T2, V5, D7, P8, E155, W156, P157, D162, T184, M191, R192, K193, P196, D258, F261, S263, N264, V267, K282, L284, D290. Residues of RrGFP: V48, A52, P53, L54, P55, F58, D59, M132, M136, I140, V141, T169, T207, S209, G210, Y211, V212. The solvent-accessible surface area (SASA) of the RrLuc−RrGFP complex formation is 7.6±0.2% for RrLuc and 8.8±0.2% for RrGFP. Solvation energy gain on complex formation (Δ^i^G) is -4.7±1.2 kcal/mol for RrLuc and -2.6±0.3 kcal/mol for RrGFP.

| RrLuc | RrGFP | Length (Å) |
| --- | --- | --- |
| H-bonds | | |
| P259 [O] | R131 [NE] | 3.01 |
| P259 [O] | R131 [NH2] | 2.97 |
| E256 [OE1] | R131 [NH1] | 3.43 |
| S257 [O] | R131 [NH2] | 3.00 |
| G260 [O] | Q137 [NE2] | 2.42 |
| S188 [OG] | Q168 [NE2] | 3.13 |
| D158 [OD2] | K171 [NZ] | 2.75 |
| S3 [OG] | K202 [NZ] | 3.19 |
| K189 [NZ] | D139 [OD1] | 3.24 |
| K189 [NZ] | Q168 [OE1] | 3.40 |
| Y6 [OH] | A208 [O] | 3.08 |
| K4 [NZ] | Q138 [OE1] | 3.32 |
| Salt bridges | | |
| E256 [OE2] | R131 [NH1] | 3.10 |
| E256 [OE2] | R131 [NH2] | 2.57 |
| D158 [OD1] | K171 [NZ] | 3.29 |

**Extended Data Table 5. Determined pre-steady-state and steady-state kinetic values and parameters for unbound and RrGFP-bound RrLuc luciferase according to the induced-fit mechanism.** The parameters were obtained by global fitting using numerical integration and by confidence contour analysis for χ^2^ threshold of 0.95. The provided rate constants correspond to values at 15 °C in case of the pre-steady-state parameters and 37 °C in case of steady-state parameters and the inhibition constants. The values of dissociation constants *K*_d_ and *K*_d,app_ were calculated as combinations of rate constants (**Eq. 7**) and their respective standard errors were determined according to the error propagation rules. The Michaelis constant *K*_m_, dissociation constants *K*_d,EP_, *K*_d,EI_ and *K*_d,EPI_, the relative specific total luminescence intensity (TLI_s,rel_) and their respective standard errors and confidence intervals were obtained from direct fitting of the steady-state data, and the value of *k*_cat_/*K*_m_ was calculated as their simple ratio. n.a. = not applicable. ^†^ = value was obtained by on-chip pre-steady-state measurement. ^‡^ = value was obtained by a conventional stopped-flow experiment.

| Parameter | RrLuc uncomplexed | | RrLuc−RrGFP complex | |
| --- | --- | --- | --- | --- |
|  | Value ± S.E. | Confidence intervals | Value ± S.E. | Confidence intervals |
| *k*_+1_ [µM^-1^.s^-1^] | 2.7 ± 0.2^†^ | 2.5–3.1^†^ | 14.0 ± 0.8^†^ | 11.2–18.1^†^ |
|  | 5.46 ± 0.03^‡^ | 5.04–5.90^‡^ | 14.1 ± 0.2^‡^ | 10.7–20.8^‡^ |
| *k*_-1_ [s^-1^] | 304 ± 16^†^ | 252–380^†^ | 157 ± 29^†^ | 64–298^†^ |
|  | 250 ± 7^‡^ | 138–503^‡^ | 140 ± 1^‡^ | 128–153^‡^ |
| *k*_+2_ [s^-1^] | 44 ± 4^†^ | 33–63^†^ | 132 ± 4^†^ | 116–152^†^ |
|  | 91.4 ± 0.4^‡^ | 84–101^‡^ | 116 ± 1^‡^ | 90–150^‡^ |
| *k*_-2_ [s^-1^] | 26 ± 3^†^ | 17–38^†^ | 15 ± 2^†^ | 6–27^†^ |
|  | 14.7 ± 0.1^‡^ | 13.7–16.1^‡^ | 6.62 ± 0.04^‡^ | 5.80–7.76^‡^ |
| *K*_d,ES_ [µM] | 113 ± 10^†^ | n.a. | 11 ± 2^†^ | n.a. |
|  | 46 ± 2^‡^ |  | 6.4 ± 0.2^‡^ |  |
| *K*_d,ES(app)_ [µM] | 42 ± 7^†^ | n.a. | 1.1 ± 0.3^†^ | n.a. |
|  | 11 ± 2^‡^ |  | 0.54 ± 0.03^‡^ |  |
| *E*_a,+1_ [kJ.mol^-1^] | 21 ± 7^†^ | 7–38^†^ | 43 ± 9^†^ | 22–77^†^ |
| *E*_a,-1_ [kJ.mol^-1^] | 74 ± 7^†^ | 56–96^†^ | < 89^†^ | 0–89^†^ |
| *E*_a,+2_ [kJ.mol^-1^] | < 36^†^ | 0–36^†^ | < 16^†^ | 0–16^†^ |
| *E*_a,-2_ [kJ.mol^-1^] | 95 ± 13^†^ | 61–133^†^ | < 105^†^ | 0–105^†^ |
| *K*_m_ [µM] | 1.8 ± 0.1 | 1.6–2.0 | 1.0 ± 0.1 | 0.9–1.2 |
| *k*_cat_ [s^-1^] | 2.76 ± 0.08 | 2.62–2.93 | 16.0 ± 0.5 | 15.2–16.8 |
| *k*_cat_/*K*_m_ [s^-1^.µM^-1^] | 1.50 ± 0.05 | n.a | 13.2 ± 1.1 | n.a. |
| *K*_d,EP_ [µM] | 1.28 ± 0.07 | 1.15–1.43 | 1.7 ± 0.2 | 1.5–2.0 |
| *K*_d,EI_ [µM] | 11.9 ± 0.1 | 11.4–12.6 | 11.1 ± 0.1 | 10.1–12.3 |
| *K*_d,EPI_ [µM] | 11.6 ± 0.1 | 11.0–12.3 | n.a. | n.a. |
| TLI_s,rel_ [%] | 100 | n.a. | 194 | n.a. |

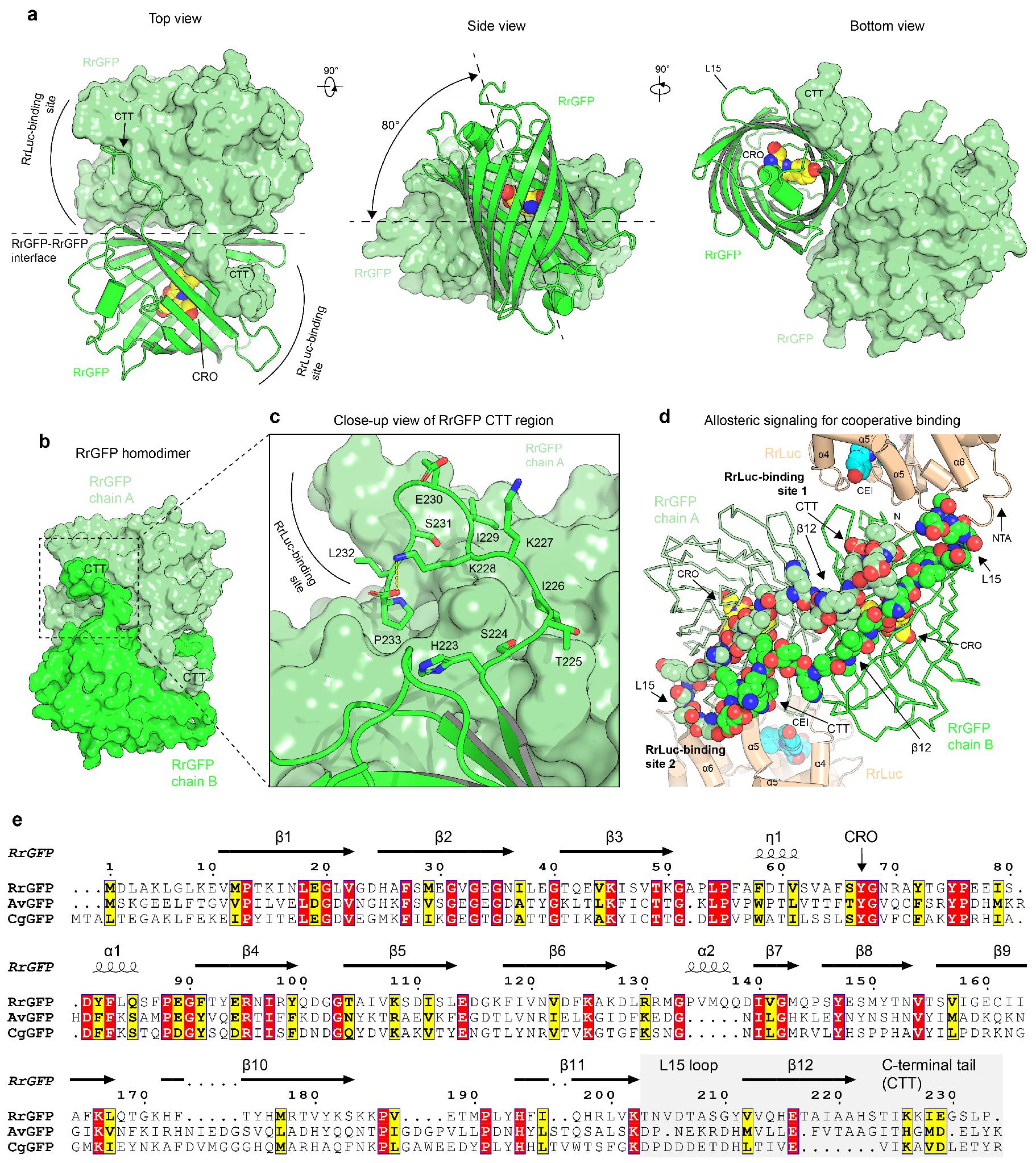

**Extended Data Fig. 1. Molecular details of the RrGFP homodimer.** (**a**) Structural views of RrGFP homodimer when bound to RrLuc luciferase. One RrGFP protomer is shown in a surface representation, while the other protomer is shown as a cartoon representation. CRO fluorophore shown as yellow spheres. (**b**) Surface representation highlighting the role of RrGFP CTT region in tight homodimerisation. Chains A and B are coloured in different shades of green. (**c**) Close-up view of RrGFP CTT region, revealing how one RrGFP protomer shields the surface of a second protomer. (**d**) Visualising a putative allosteric communication pathway in the *Renilla* BRET complex. Ribbon representation of RrGFP homodimer (shades of green) with two RrLuc luciferase molecules bound (beige). Structural elements (L15, β12, and CTT), which connect the RrLuc-binding site 1 and RrLuc-binding site 2, are shown in a space-filling spheres representation. (**e**) Sequence alignment of GFP proteins from *R*. *reniformis* (RrGFP), *Aequorea victoria* (AvGFP), and *Clytia gregaria* (CgGFP). The structural elements that constitute the putative allosteric communication pathway (L15, β12, and CTT) are highlighted in a shade of grey. The topology of secondary structure elements is shown over the alignment.

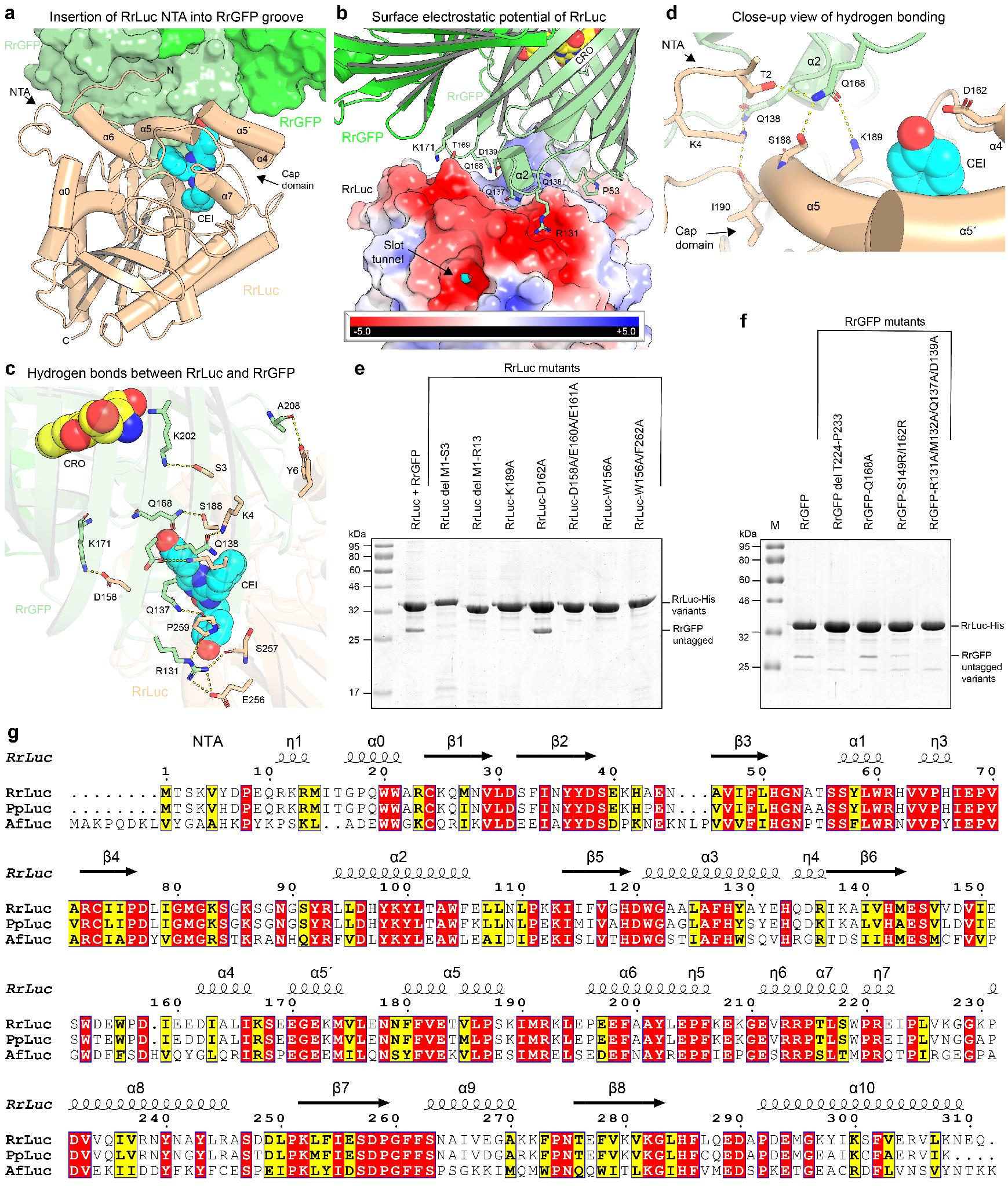

**Extended Data Fig. 2. Molecular features of the RrGFP−RrLuc complexation.** (**a**) Structural snapshot that shows the insertion of RrLuc N-terminal arm (NTA) into a groove shaped on the RrGFP homodimer surface. Note that the RrLuc cap domain (helices α4, α5´ and α5) physically interacts with the RrGFP. CEI emitter shown as cyan spheres. (**b**) Protein-protein interactions between RrLuc and RrGFP. Electrostatic surface potential mapped on RrLuc. The RrGFP homodimer is shown in a cartoon representation. Selected RrGFP surface residues making contacts with RrLuc are shown as green sticks. (**c**) Close-up view of the hydrogen bonding network between RrLuc (NTA and cap domain) and RrGFP. CEI emitter shown as cyan spheres. (**d**) Additional hydrogen bonding between RrLuc and RrGFP. CEI emitter shown as cyan spheres; CRO fluorophore shown as yellow spheres. (**e,f**) Mutagenesis and co-expression assays validating the crystallographic observations. Structure-based mutagenesis of RrLuc (e) and RrGFP (f) and validation of their effects on the RrGFP−RrLuc complexation via co-expression assays in *E*. *coli*. (**g**) Sequence alignment of luciferases from *R*. *reniformis* (RrLuc), *Pennatula phosphorea* (PpLuc) and *Amphiura filiformis* (AfLuc). The topology of secondary structure elements is shown over the alignment.

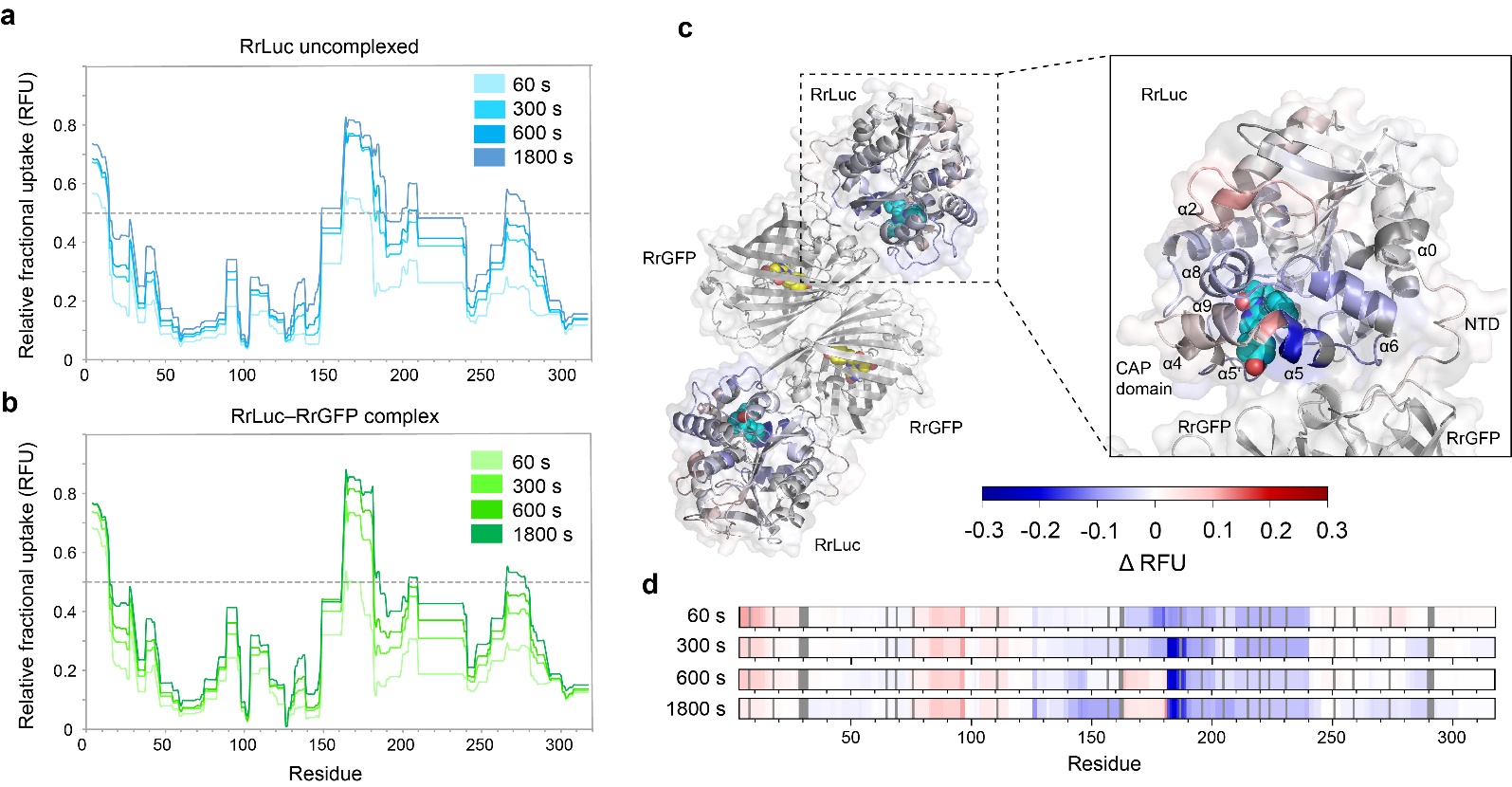

**Extended Data Fig. 3. Time-course hydrogen-deuterium exchange (HDX) mapped onto the RrLuc luciferase. (a,b)** Relative fractional uptake (RFU) of uncomplexed RrLuc luciferase (**a**) and RrGFP-bound RrLuc complex (**b**) plotted against residue number for four labelling times (60, 300, 600, and 1800 s). The topology of secondary structure elements is shown in **Extended Data Fig. 2g**. The dashed horizontal line marks 50% relative uptake. **(c)** ΔRFU (ΔRFU = RFU of RrLuc-RrGFP complex − RFU of unbound RrLuc) after 1800 s labelling time mapped onto the structure of the RrLuc–RrGFP complex, PDB ID: 8RZZ. Blue indicates regions that are protected upon complex formation, exchanging less than in the free form; red indicates regions that exchange more upon RrGFP binding. RrGFP subunits (grey) are shown for context only and were not analysed. Note that transient differences in deuterium uptake at the N-terminal arm, most pronounced at early labelling times (see panel **d**), are largely resolved by 1800 s and are therefore not prominent in this view. **(d)** ΔRFU heat maps mapped onto the RrLuc sequence at each of the four labelling times (60, 300, 600, and 1800 s, top to bottom), using the same ΔRFU convention as in panel (**c**). Blue regions indicate protection upon complex formation (decreased amide hydrogen (H) to deuterium (D) exchange relative to the unbound RrLuc protein), while red regions indicate increased exchange upon binding. Proline residues, non-exchangeable in HDX-MS, and unresolved residues are indicated by dark and light grey rectangles, respectively.

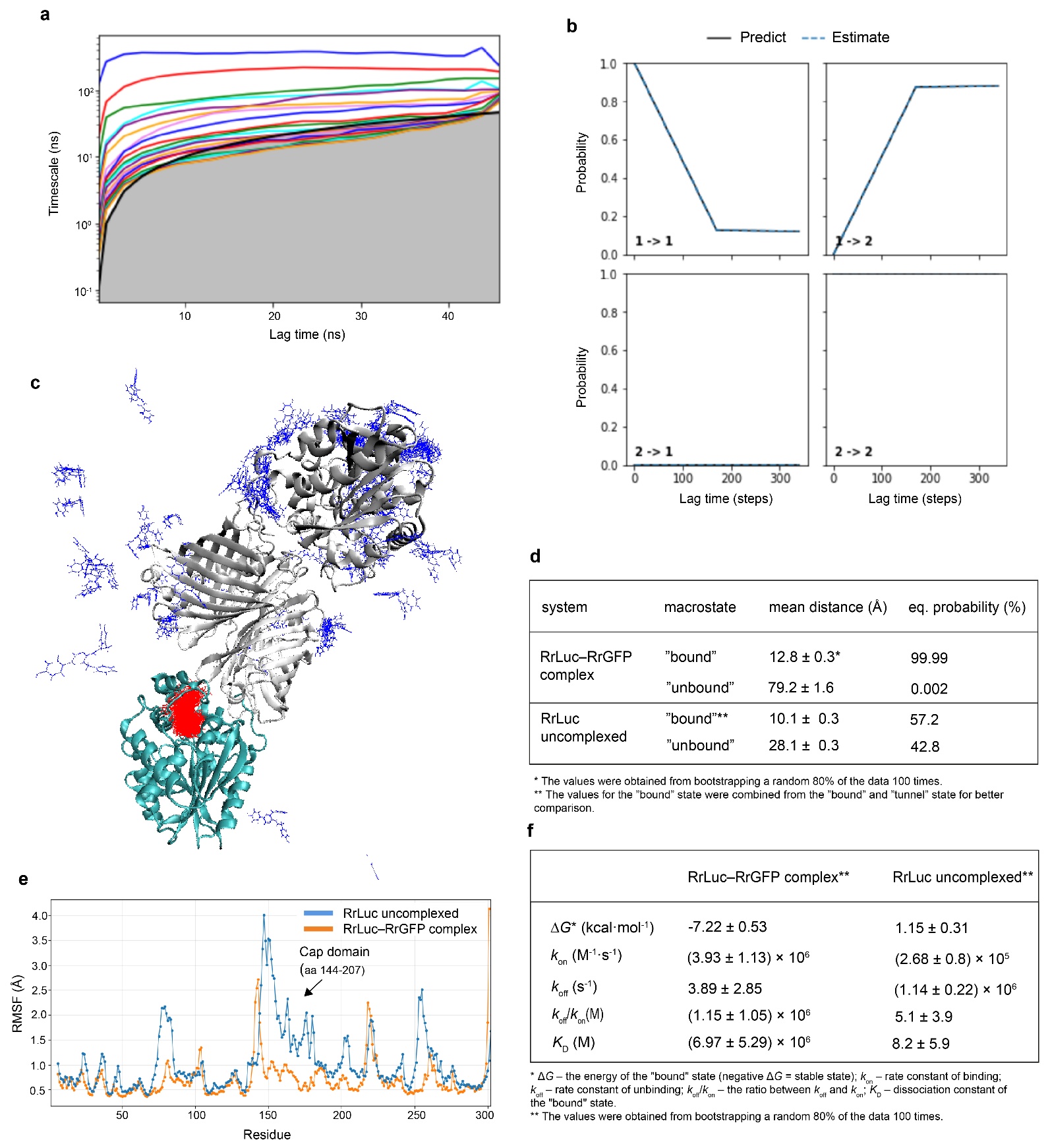

**Extended Data Fig. 4.** **Protein simulations.** (**a**,**b**) Identification and quality control of Markov states. Implied timescales and Chapman-Kolmogorov test of MSM of CEI release from the RrLuc–RrGRP complex system. The implied timescales (a) show the transition between the macrostates. The Chapman-Kolmogorov test (b) was done for a Markov state model of three macrostates constructed at a 17 ns lag time. The selected lag time is the lowest at which the “predict” (black) and “estimate” (blue dashes) lines agree. **(c)** CEI macrostates from Markov state model (MSM) from adaptive sampling of CEI unbinding from RrLuc (cyan cartoon) within the RrLuc–RrGFP complex (gray cartoon). The macrostates, defined by the distance of the ligand from the active site, are shown as thin sticks and coloured based on their distance to the active site (“bound” state in red and “unbound” state in blue). (**d**,**e**) Dynamics of the uncomplexed versus the RrGFP-bound RrLuc luciferase. (**d**) Summary of the mean CEI-active site distance and the equilibrium probabilities of macrostates from the adaptive sampling of CEI release from uncomplexed and RrGRP-complexed RrLuc. The equilibrium probability shifts towards the bound form in the complexed form, implying altered product release dynamics. **(e)** Root-mean square flexibility (RMSF) of uncomplexed RrLuc (blue) and the RrGFP-bound RrLuc (orange). Global profiles are maintained with locally higher flexibility for uncomplexed RrLuc. However, in the cap domain, flexibility is notably decreased in RrLuc. (**f**) The CEI unbinding kinetics from adaptive sampling simulations with the uncomplexed RrLuc and the RrGFP-bound RrLuc luciferase. The kinetic parameters were calculated between the “bound” and “unbound” states.

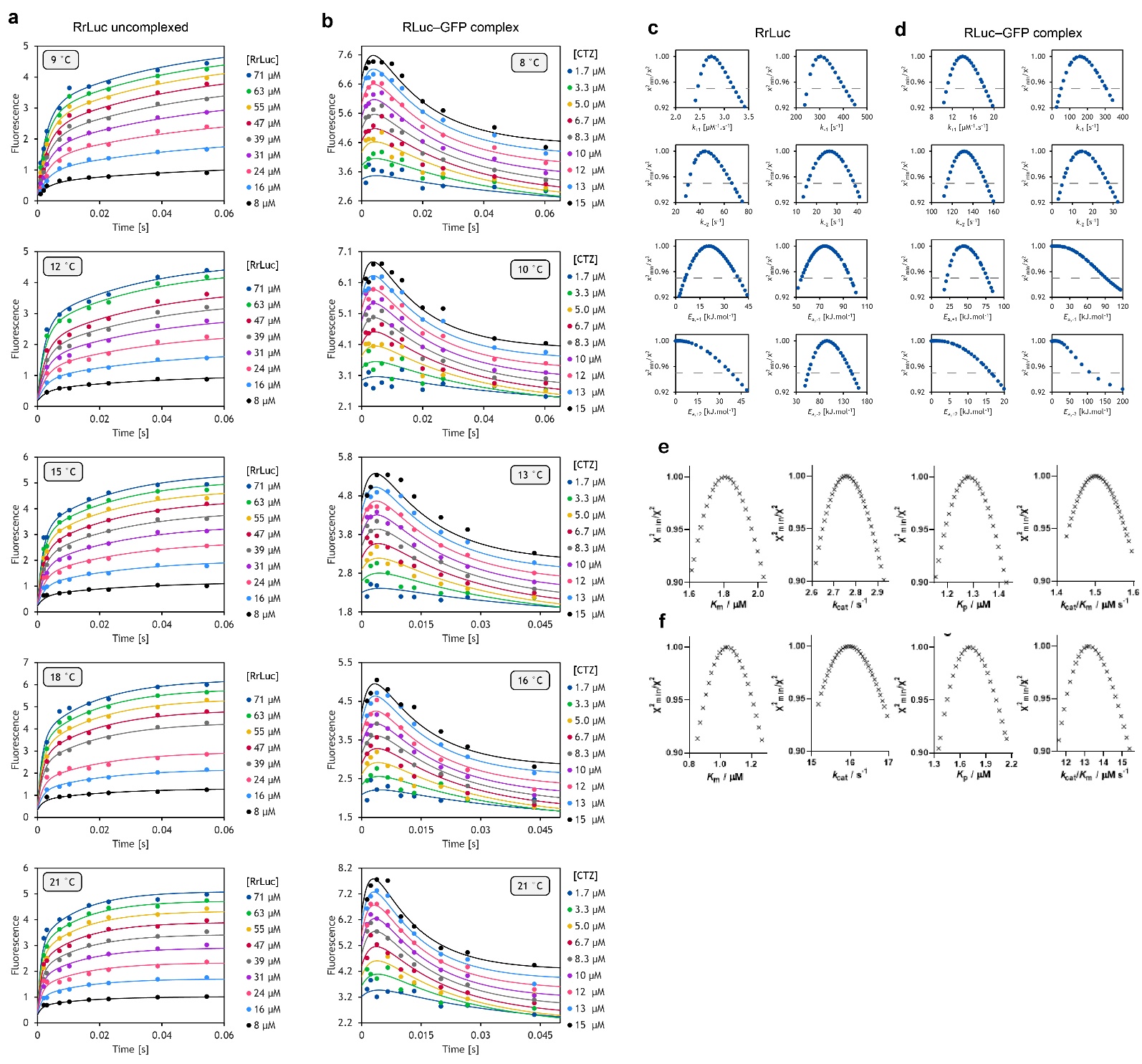

**Extended Data Fig. 5. Global numerical fitting and confidence contour analyses of kinetic data.** (**a**,**b**) Global numerical fitting of the CTZ fluorescence kinetic curves upon the CTZ binding at various temperatures by either uncomplexed RrLuc (**a**) or RrLuc−RrGFP complex (**b**). The data were fitted using the induced-fit binding mechanism (**Fig. 6e**, **Eq. 5**) for both the analysed systems. Solid lines represent the best fit. The data were collected in 5 mM KCl, 10 mM Tris-HCl buffer (pH 7.5) at temperatures specified in each graph. Every datapoint corresponds to an average of three independent technical replicates. (**c**,**d**) Comparison of confidence contour analyses of parameters used for fitting kinetic data of either uncomplexed RrLuc (**c**) or RrGFP-complexed RrLuc (**d**) according to the induced-fit mechanism. Lower bound limits of several activation energies (*E*_a,-1_, *E*_a,+2_ , *E*_a,-2_) were not well defined by the data, allowing us to assess only their upper limits. All the remaining parameters were well constrained by the collected kinetic data, confirming the robustness of the performed analysis. The grey dashed line represents the χ^2^ threshold of 0.95. (**e**,**f**) 1D confidence contour analysis of RrLuc catalytic parameters. The upper panels correspond to the uncomplexed RrLuc (**e**) and the bottom ones to the RrGFP-bound RrLuc (**f**).
